# Functional characterization of the language network of polyglots and hyperpolyglots with precision fMRI

**DOI:** 10.1101/2023.01.19.524657

**Authors:** Saima Malik-Moraleda, Olessia Jouravlev, Maya Taliaferro, Zachary Mineroff, Theodore Cucu, Kyle Mahowald, Idan A. Blank, Evelina Fedorenko

## Abstract

How do polyglots—individuals who speak five or more languages—process their languages, and what can this population tell us about the language system? Using fMRI, we identified the language network in each of 34 polyglots (including 16 hyperpolyglots with knowledge of 10+ languages) and examined its response to the native language, non-native languages of varying proficiency, and unfamiliar languages. All language conditions engaged all areas of the language network relative to a control condition. Languages that participants rated as higher-proficiency elicited stronger responses, except for the native language, which elicited a similar or lower response than a non-native language of similar proficiency. Furthermore, unfamiliar languages that were typologically related to the participants’ high-to-moderate-proficiency languages elicited a stronger response than unfamiliar unrelated languages. The results suggest that the language network’s response magnitude scales with the degree of engagement of linguistic computations (e.g., related to lexical access and syntactic-structure building). We also replicated a prior finding of weaker responses to native language in polyglots than non-polyglot bilinguals. These results contribute to our understanding of how multiple languages co-exist within a single brain and provide new evidence that the language network responds more strongly to stimuli that more fully engage linguistic computations.

## Introduction

A large fraction of the world’s population speak or sign more than one language. Not surprisingly, research in psycholinguistics and cognitive neuroscience has long tackled questions about how bilingual minds and brains work. The research questions have ranged from how multiple languages are represented and processed (e.g. Chee et al., 1999, 2000; Fabbro, 2001; Hernandez et al., 2001; Lucas et al., 2004; Perani & Abutalebi, 2005; Liu & Cao, 2016), to whether native language processing differs between bilingual and monolingual individuals (e.g., Kovelman et al., 2008; Jones et al., 2012; Grundy et al., 2017; Pliatsikas et al., 2020; Arredondo et al., 2022), to whether bilingualism confers advantages outside of the language domain (e.g., Bialystok et al., 2016; Cespón & Carreiras, 2020; Blanco-Elorrieta & Caramazza, 2021).

Although a handful of studies have investigated individuals who speak three or more languages (e.g., Vingerhoets et al., 2003; Briellmann et al., 2004; Jeong et al., 2007; Lemhöfer et al., 2010; Videsott et al., 2010; Abutalebi et al., 2013a,b; De Bruin et al., 2014; Kong et al., 2014; Yang et al., 2018; Yazbek et al., 2020), the bulk of past work has focused on bilinguals. However, for some research questions, bilinguals are not an ideal test population. First, by definition, research on bilinguals is limited to asking questions about two languages, and some findings about the representation and processing of two languages may not generalize to the representation and processing of multiple languages. For example, although many have argued that in bilinguals, the two languages draw on the same brain areas (e.g., Illes et al., 1999; Roux & Trétmoulet, 2002; Klein et al., 2006; Sulpizio et al., 2020), it is possible that additional languages (perhaps the later-acquired ones or the lower-proficiency ones) would not show the same pattern and would instead draw on a system that supports the acquisition of novel cognitive skills later in life (e.g., Abutalebi, 2008; Liu et al., 2010; Yazbek et al., 2020). And second, in many cases, comparisons between a bilingual’s two languages involve a comparison between a native language (L1) and a second, non-native language acquired later in life (L2). Because one’s native language may have a privileged status in how it is represented and processed (e.g., Cutler, 2012; Keysar et al., 2012; Pierce et al., 2014), such comparisons may be difficult to interpret when trying to understand, for example, how proficiency influences neural responses (e.g., is L2 eliciting a weaker/stronger response than L1 because the individual is less proficient in it or because it is not the individual’s native language?). Examining neural responses to several non-native languages can therefore help clarify how different aspects of one’s experience with a language (here, we focus on proficiency specifically) affect brain responses.

Furthermore, past cognitive neuroscience research on bilingualism has suffered from limitations. First, much past work has relied on the traditional group-averaging fMRI approach (e.g., comparing group-level activation maps for L1 vs. L2), which suffers from low sensitivity, low functional resolution, and low interpretability (e.g., Saxe et al., 2006; Nieto-Castañón & Fedorenko, 2012; Fedorenko, 2021). And second, many studies have relied on paradigms that conflate language processing and general task demands, which recruit distinct brain networks: the language-selective network (Fedorenko et al., 2011) and the domain-general Multiple Demand network (e.g., Duncan, 2010, 2013), respectively (see Fedorenko & Blank, 2022 for review). As a result, overlap between languages may reflect similar task demands in addition to, or even instead of, language representation/processing.

To address these limitations, we here i) turn to polyglots and hyperpolyglots, who have some degree of proficiency in at least 5 languages (range in our sample: 5-54 languages), and ii) use robust individual-level fMRI analyses and an extensively validated language network ‘localizer’ paradigm (e.g., Fedorenko et al., 2010; Braga et al., 2020; Lipkin et al., 2022; Malik-Moraleda, Ayyash et al., 2022). We ask two questions that have so far been mostly probed in bilinguals and that remain debated (e.g., Costa & Sebastián-Gallés, 2014; Sulpizio et al., 2020; Pascual et al., 2023). First, we ask whether multiple languages (including the native language, three other languages of varying proficiency, and unfamiliar languages) all draw on the fronto-temporal language-selective network. And second, we ask how proficiency affects the magnitude of neural response in the language areas. In addition, we ask a novel question about neural responses to unfamiliar languages that are typologically related to the participants’ high-/moderate-proficiency languages. Given that, due to a common ancestral language, related languages (sometimes referred to as ‘sister languages’; e.g., Campbell, 2017) exhibit overlap in their sound patterns, vocabulary, and grammatical constructions, participants may be able to extract some meaning from those languages (e.g., Gooskens et al., 2017) or at least to recognize familiar sound patterns. We test whether this (relatively minimal) level of comprehension would manifest as a stronger response in the language areas compared to completely unfamiliar unrelated languages, which should be incomprehensible. Finally, in line with increasing emphasis on robustness and replicability in cognitive neuroscience (e.g., Poldrack et al. 2017), we attempt to replicate a recent finding of lower neural responses during native language processing in polyglots compared to non-polyglots (Jouravlev et al., 2021).

To foreshadow our key results, all languages, including completely unfamiliar ones, engage the entire fronto-temporal language-selective brain network to a greater degree than a perceptually-matched control condition. The level of response to different languages generally scales with self-reported proficiency. This pattern of results plausibly reflects a greater ability of higher-proficiency languages to engage in linguistic computations, although greater co-activation of the native language during the processing of higher-proficiency languages may also play a role.

## Methods

### Participants

Candidate (self-proclaimed) polyglots were recruited by word of mouth. Most participants resided in the Boston area, but a few traveled from other cities in the U.S. Candidate participants were asked to assess their proficiency in each language that they had some familiarity with by rating their listening, speaking, reading, and writing abilities on a scale from 0=no ability to 5=native/native-like ability. The four scores were summed to obtain an overall proficiency score for each language, which could range from 1 (minimal proficiency) to 20 (native/native-like proficiency) (see Malik-Moraleda et al., 2023, for evidence—from a different population—that this subjective measure correlates with more objective proficiency measures). Similar to Jouravlev et al. (2021), a polyglot was defined as an individual who self-reported (a) at least minimal proficiency (a score of 4 out of 20) in at least 5 languages (i.e., their native language and four other languages; for five participants, proficiency ratings were only collected for the 4 familiar languages tested but they did informally report some knowledge of five or more languages, based on which they were recruited for the study; **Table S1**), and (b) moderate-to-high proficiency (an overall score of at least 10 out of 20) in at least one language other than their native language (all but three participants had a score of 16 or higher on their second most proficient language, and two of these three had a score of 10 or higher on their third most proficient language). Based on these criteria, 35 participants were included in the study. One participant’s data were excluded due to excessive in-scanner motion, leaving 34 polyglot participants for analysis.

Participants were between 19 and 71 years of age at the time of testing (*M*=35.0 years; *SD*=13.9). Most (21 of the 34; ∼60%) were native speakers of English; the remaining thirteen were native speakers of French (n=4), Russian (n=3), Spanish (n=2), Dutch (n=1), German (n=1), Hungarian (n=1), and Mandarin (n=1) and proficient speakers of English (proficiency range for English across these 13 participants: 14-20, *M*=19.2, *SD*=1.69). Twenty participants (∼59%) were male, and 31 (∼91%) were right-handed.

The mean number of languages spoken/signed was 14.6 (median=10, range: 5-54 languages; **Table S1**). The mean self-reported proficiency for the highest-proficiency language was 20 out of the maximum score of 20 (*SD*=0; note that for three participants, the highest-proficiency language was not their native language; **Table S1**), 18.3 for the second most proficient language (*SD*=2.45, range: 10-20), 15.8 for the third most proficient language (*SD*=3.36, range: 8-20), 13.3 for the fourth most proficient language (*SD*=4.034, range: 7-20), 11.2 for the fifth most proficient language (*SD*=4.23, range: 4-20), and—for individuals with proficiency in more than five languages—10.33 for the sixth most proficient language (*SD*=3.99, range: 4-18). Thus, in addition to having native-like proficiency in their second most proficient language, many participants had high proficiency in their third and fourth most proficient languages, and some in their fifth and sixth most proficient languages (**Table S1**).

All but one participant showed typical, left-lateralized language activations during the language localizer task (described below), as assessed by a lateralization index (LI) (e.g., Jouravlev & Jared, 2020). In particular, we i) calculated the number of voxels—within the boundaries of the language parcels in the left hemisphere (LH) and the right hemisphere (RH) (see <u>Language fROI definition and response estimation</u> below)—that are significant for the language localizer contrast at a fixed (p<0.001 uncorrected whole-brain) statistical threshold, and then ii) used the following formula “LI = (number of LH voxels - number of RH voxels) / (number of LH voxels + number of RH voxels)”. The LI values range between -1 (exclusively right-hemisphere activations) to +1 (exclusively left-hemisphere activations). The mean LI was 0.429 (*SD*=0.363; range: -0.850-0.930); the mean LI for the 33 participants, when excluding the right-lateralized participant, was 0.468 (*SD*=0.288; range: -0.181-0.93; **Table S2**).

For one analysis, we additionally used a previously published dataset of eighty-six non-polyglot bilingual individuals, who were native speakers of diverse languages and had completed a language listening task in their native language (Malik-Moraleda, Ayyash et al., 2022), similar to the language listening task in the current study. Participants were between 19 and 45 years of age at the time of testing (*M*=27.5, *SD*=5.49). Half of the participants (n=43) were male. All participants were right-handed. The mean LI was 0.586 (*SD*=0.292; range: -0.180-0.997).

All participants gave informed written consent in accordance with the requirements of the Committee on the Use of Humans as Experimental Subjects at MIT and were paid for their participation.

## Experimental Design and Materials

Each participant completed a localizer for the language network (Fedorenko et al., 2010) and the critical multi-language listening task. Some participants completed one or two additional tasks for unrelated studies. The entire scanning session lasted approximately two hours.

### Language localizer

Participants passively read sentences and lists of pronounceable nonwords in a blocked design. The *Sentences*>*Nonwords* contrast targets brain regions that are sensitive to high-level linguistic processing, including the understanding of word meanings and combinatorial phrase-structure building (Fedorenko et al., 2010; Shain, Kean et al., 2023; see Fedorenko et al., in press, for review). In prior work, we had established the robustness of the language localizer to changes in the materials, modality of presentation (reading vs. listening), task, timing parameters, and other aspects of the procedure (e.g., Fedorenko et al., 2010; Fedorenko, 2014; Mahowald & Fedorenko, 2016; Scott et al., 2017; Cheung et al., 2020; Lipkin et al., 2022; Malik-Moraleda, Ayyash et al., 2022). We used an English version of this localizer for all participants (see Malik-Moraleda, Ayyash et al., 2022 for evidence that this localizer works well for non-native proficient speakers of English, identifying the same areas as a localizer contrast based on a participant’s native language). Each trial started with 100 ms pre-trial fixation, followed by a 12-word-long sentence or a list of 12 nonwords presented on the screen one word/nonword at a time at the rate of 450 ms per word/nonword. Then, a line drawing of a hand pressing a button appeared for 400 ms, and participants were instructed to press a button whenever they saw this icon, and finally a blank screen was shown for 100 ms, for a total trial duration of 6 s. The simple button-pressing task was included to help participants stay awake and focused. Each block consisted of 3 trials and lasted 18 s. Each run consisted of 16 experimental blocks (8 per condition), and five fixation blocks (14 s each), for a total duration of 358 s (5 min 58 s). Each participant performed two runs. Condition order was counterbalanced across runs.

### Multi-language listening task

Participants listened to brief passages in eight languages and to control, acoustically scrambled versions of those passages (see below for details) in a blocked design. The eight languages were selected separately for each participant based on their linguistic background (**Tables S1, S3**) and included (a) the participant’s ***native*** language (L1), (b) three ***non-native*** languages that the participant was somewhat proficient in (L2, L3, and L4; see below for proficiency details), (c) two unfamiliar languages that were ***related*** to one or more of the participant’s high-to-moderate proficiency languages (**Tables S3, S6**; the proficiency level for the language to which an unfamiliar language was related was: *M=*17.9*, SD=*3.36, range: 8-20), and two languages that the participant was completely ***unfamiliar*** with and that were unrelated to any of the participant’s languages (**Tables S3, S6**). For the languages used in the experiment, L1 was the participant’s native language, with the mean self-rated proficiency of 19.4 out of the maximum score of 20 (*SD=*2.16, range: 10-20; all but three participants rated themselves to have a proficiency of 20). L2 was the non-native language that participants reported being most proficient in, with the mean proficiency of 18.7 out of 20 (*SD*=1.82, range: 12-20) (for the 3 participants whose native language was not the most proficient language, their L2 was their most proficient language, with proficiency of 20). L3 and L4 were chosen so that participants were somewhat proficient in them (a score of at least 4 out of 20 in each), but these were not always the next most proficient languages due to the limitations on the languages for which experimental materials were available. The mean proficiency for L3 was 13.5 (*SD*=3.34, range: 6-20) and for L4, it was 9.15 (*SD*=3.29, range: 4-16). Across participants, materials from 34 languages were used (**Table S3**).

Two sets of materials were used in the multi-language listening experiment. One set (used for n=18 participants) came from the publicly available corpus of Bible audio stories (https://www.biblegateway.com/resources/audio/), which consists of Bible-based stories that are narrated by native speakers of different languages. This set included materials for 25 languages (Arabic, Bangla, Basque, Dutch, English, French, Georgian, German, Hausa, Hebrew, Hindi, Indonesian, Italian, Japanese, Mandarin, Romanian, Russian, Persian, Spanish, Swahili, Thai, Turkish, Wolof, Xhosa, and Yiddish), 18 of which were used in the current study. The other set (used for the remaining n=16 participants) used passages from *Alice in Wonderland* (Carroll, 1865). The passages are narrated by native speakers, and the recordings were created in the lab for an earlier study (Malik-Moraleda, Ayyash et al., 2022) (https://evlab.mit.edu/aliceloc/). This set included materials for 46 languages (Afrikaans, Arabic, Armenian, Assamese, Basque, Belarusian, Bulgarian, Catalan, Czech, Danish, Dutch, English, Farsi, Finnish, French, German, Greek, Gujarati, Hebrew, Hindi, Hungarian, Irish, Italian, Japanese, Korean, Latvian, Lithuanian, Mandarin, Marathi, Nepali, Norwegian, Polish, Portuguese, Romanian, Russian, Serbo-Croatian, Slovene, Spanish, Swahili, Swedish, Tagalog, Tamil, Telugu, Turkish, Ukrainian, and Vietnamese), 28 of which were used in the current study. The switch to the Alice in Wonderland materials was made because of the larger number of languages available in that set and the fact that new languages are continually being added to it. (Although in all the main analyses, we analyze all 34 participants together, we report the results for the two sets of materials separately in **Figures S3, S5c**).

For both the Bible stories and Alice in Wonderland passages, for each critical language condition, a set of 8 audio clips was selected, each 16 s long. The control condition clips were created using a sound “quilting” procedure (Overath et al., 2015). In particular, for each language, a 1-1.5 min clip of continuous speech was selected (from the respective corpus: the Bible stories corpus or the Alice in Wonderland materials; for the Bible stories, the clips were selected from the available materials for each language, for the Alice materials, one of the long passages was used—see Malik-Moraleda, Ayyash et al., 2022 for details). These clips were divided into 30 ms segments, and the segments were then re-ordered in such a way that (a) the segments adjacent to each other in the original clip are not adjacent, (b) segment-to-segment cochleogram changes match the original signal as closely as possible, and (c) boundary artifacts are minimized. Eight quilted clips, each 16 s long, were created for each language. Finally, for all the resulting intact and quilted clips, sound intensity was normalized, and each clip was further edited to include brief (1 s long) fade-in and fade-out periods at the beginning and end, respectively. Intensity normalization and fade-in/fade-out editing were performed using the Audacity software (Audacity, 2014) for the Bible stories materials and Matlab (Mathworks, 2020) for the Alice in Wonderland materials. All the materials used in this experiment are available at: https://osf.io/3he75/. The experimental script is available upon request. For each set of materials (Bible stories and Alice in Wonderland passages), the full set (64 intact and 64 quilted clips) was divided into 8 subsets, which corresponded to 8 scanning runs. Each run consisted of 8 intact clips (1 clip per language) and 8 quilted clips (1 clip per language) and 3 fixation periods each lasting 12 s, for a total run duration of 292 s (4 min 52 s). Each participant performed 8 runs. The order of conditions (intact, quilted) and languages was counterbalanced across runs and participants.

### fMRI data acquisition

Structural and functional data were collected on the whole-body 3 Tesla Siemens Trio scanner with a 32-channel head coil at the Athinoula A. Martinos Imaging Center at the McGovern Institute for Brain Research at MIT. T1-weighted structural images were collected in 179 sagittal slices with 1 mm isotropic voxels (TR=2,530 ms, TE=3.48 ms). Functional, blood oxygenation level dependent (BOLD) data were acquired using an EPI sequence (with a 90° flip angle and using GRAPPA with an acceleration factor of 2), with the following acquisition parameters: thirty-one 4mm thick near-axial slices, acquired in an interleaved order with a 10% distance factor; 2.1 mm x 2.1 mm in-plane resolution; field of view of 200 mm in the phase encoding anterior to posterior (A>P) direction; matrix size of 96 x 96 voxels; TR of 2,000 ms; and TE of 30 ms. Prospective acquisition correction (Thesen et al., 2000) was used to adjust the positions of the gradients based on the participant’s motion one TR back. The first 10 s of each run were excluded to allow for steady-state magnetization.

### fMRI data preprocessing

fMRI data were analyzed using SPM12 (release 7487), CONN EvLab module (release 19b), and other custom MATLAB scripts. Each participant’s functional and structural data were converted from DICOM to NIFTI format. All functional scans were coregistered and resampled using B-spline interpolation to the first scan of the first session (Friston et al., 1995). Potential outlier scans were identified from the resulting subject-motion estimates as well as from BOLD signal indicators using default thresholds in CONN preprocessing pipeline (5 standard deviations above the mean in global BOLD signal change, or framewise displacement values above 0.9 mm; Nieto-Castañón, 2020). Functional and structural data were independently normalized into a common space (the Montreal Neurological Institute [MNI] template; IXI549Space) using SPM12 unified segmentation and normalization procedure (Ashburner & Friston, 2005) with a reference functional image computed as the mean functional data after realignment across all timepoints omitting outlier scans. The output data were resampled to a common bounding box between MNI-space coordinates (-90, -126, -72) and (90, 90, 108), using 2 mm isotropic voxels and 4th order spline interpolation for the functional data, and 1 mm isotropic voxels and trilinear interpolation for the structural data. Last, the functional data were smoothed spatially using spatial convolution with a 4 mm FWHM Gaussian kernel.

### First-level modeling

Effects were estimated using a General Linear Model (GLM) in which each experimental condition was modeled with a boxcar function convolved with the canonical hemodynamic response function (HRF) (fixation was modeled implicitly, such that all timepoints that did not correspond to one of the conditions were assumed to correspond to a fixation period). Temporal autocorrelations in the BOLD signal timeseries were accounted for by a combination of high-pass filtering with a 128 s cutoff, and whitening using an AR(0.2) model (first-order autoregressive model linearized around the coefficient a=0.2) to approximate the observed covariance of the functional data in the context of Restricted Maximum Likelihood estimation (ReML). In addition to experimental condition effects, the GLM design included first-order temporal derivatives for each condition (included to model variability in the HRF delays), as well as nuisance regressors to control for the effect of slow linear drifts, subject-motion parameters, and potential outlier scans on the BOLD signal.

### Language fROI definition and response estimation

For each participant, functional regions of interest (fROIs) were defined using the Group-constrained Subject-Specific (GSS) approach (Fedorenko et al., 2010), whereby a set of *parcels or “search spaces”* (i.e., brain areas within which most individuals in prior studies showed activity for the localizer contrast) is combined with *each individual participant’s activation map* for the same or similar contrast.

To define the language fROIs, we used five parcels derived from a group-level representation of data for the *Sentences*>*Nonwords* contrast in 220 independent participants (**Figure 2a**). These parcels were used in much prior work (e.g., Pereira et al., 2018; Fedorenko et al., 2020; Malik-Moraleda, Ayyash et al., 2022; Hu, Small et al., 2022) and included three regions in the left frontal cortex: two located in the inferior frontal gyrus (LIFG and LIFGorb), and one located in the middle frontal gyrus (LMFG); and two regions in the left temporal cortex spanning the entire extent of the lateral temporal lobe (LAntTemp and LPostTemp). The fROIs were defined in each individual participant separately by selecting within each parcel the 10% of most localizer-responsive voxels based on the *t-*values for the *Sentences>Nonwords* contrast in the relevant participant’s activation map. The responses of these individually-defined fROIs to the 16 conditions of the critical task were then extracted, averaging across the voxels in each fROI. The responses to the eight quilted control conditions were further averaged in order to obtain a more robust control-condition baseline.

For completeness, we examined responses in the right hemisphere (RH) homotopes of the language regions. To define the fROIs in the RH, the left hemisphere parcels were mirror-projected onto the RH to create five homotopic parcels. By design, the parcels cover relatively large swaths of cortex in order to be able to accommodate inter-individual variability in the precise locations of functional areas. Hence the mirrored parcels are likely to encompass RH language regions despite possible hemispheric asymmetries in the precise locations of activations (for validation, see Blank et al., 2014; Mahowald & Fedorenko, 2016; Lipkin et al., 2022; Shain, Paunov, Chen et al., 2022).

## Statistical Analyses

### How does the polyglots’ language network respond to different languages?

To evaluate whether different languages all engage the language network (see <u>Methods</u>) in polyglot individuals, we examined the responses to eight languages (selected separately for each participant; see <u>Methods</u>), including four languages in which the individual had some proficiency (L1, L2, L3 and L4) as well as four languages that were unfamiliar to the participant. To do so, for each language condition, we fitted a model that predicted the BOLD response of the LH language network to that language condition with participants and fROIs modeled as random intercepts. Dummy coding was used, with the quilted control condition used as the reference level:

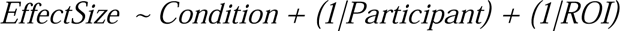

In most analyses, we treat the language network as an integrated whole. This choice is motivated by two sets of findings. First, the five language regions exhibit highly similar functional response profiles: they all show strong selectivity for language processing relative to diverse non-linguistic inputs and tasks (e.g., Fedorenko et al., 2011; Monti et al., 2012; Deen et al., 2015; Pritchett et al., 2018; Jouravlev et al., 2019; Amalric & Dehaene, 2019; Ivanova et al., 2020; Liu et al., 2020, Chen et al., 2023; for reviews, see Fedorenko & Blank, 2020 and Fedorenko et al., subm.), and they are all similarly modulated by linguistic manipulations, showing sensitivity to regularities at different levels of linguistic structure and engagement in computations related to lexical access, syntactic-structure building, and semantic composition (e.g., Fedorenko et al., 2010, 2016, 2020; Blank et al., 2016; Bautista & Wilson, 2016; Shain, Blank et al., 2020; Hu, Small et al., 2022; Shain, Kean et al., 2023; Tuckute et al., 2024). And second, these regions exhibit strong correlations in their activity during naturalistic cognition paradigms (e.g., Blank et al., 2014; Paunov et al., 2019; Malik-Moraleda, Ayyash et al., 2020; Braga et al., 2020). However, for some analyses, we additionally fitted models that predicted the BOLD response in each language fROI separately in order to explore potential differences between fROIs (see **Table S4** for the output of these LMEs; see **Figure S3** for responses in individual fROIs).

### How does self-reported proficiency affect the magnitude of response in the polyglots’ language network?

To compare the magnitudes of response in the language network to the four languages for which the participants had different proficiencies (L1-L4), we fitted a model that predicted the BOLD response in the language network with Language (L1, L2, L3, L4) modeled as a fixed effect and participants and fROIs modeled as random intercepts. Effect coding was used for the contrasts in order to make three comparisons: L1 vs. L2, L2 vs. L3, and L3 vs. L4:

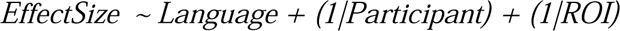

### How does the polyglots’ language network respond to unfamiliar related vs. unfamiliar unrelated languages?

Here, we focused on the comparison of unfamiliar languages that are related to one (or more) of the high-to-moderate proficiency languages of a polyglot (unfamiliar related) vs. those that are not related to any high-to-moderate proficiency languages (unfamiliar unrelated). To test whether the processing of *unfamiliar related* languages elicits a stronger response than *unfamiliar unrelated* languages (while eliciting a lower response than high-to-moderate proficiency languages), we fitted a model that predicted the BOLD response in the language network with Condition (Familiar, Related, Unrelated) modeled as a fixed effect and participants and fROIs modeled as random intercepts. Dummy coding was used, with the Related condition used as the reference level:

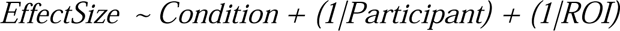

### Does the polyglots’ language network respond less strongly to native language than the language network of non-polyglots?

To our knowledge, the only past functional brain imaging investigation that focused on polyglots is Jouravlev et al. (2021) (cf. Amunts et al., 2004; Hervais-Adelman et al., 2018; Palmann & Golestani, 2020 for past *anatomical* investigations of polyglot brains; see Hervais-Adelman et al., 2015 for an fMRI study that included some polyglot participants). Jouravlev and colleagues reported that the language network of polyglots responds less strongly during native language processing compared to both a set of pairwise-matched controls and a larger group or control participants. In their study, which used a subset of the individuals tested in the current study (13 of the 34; ∼38%), Jouravlev and colleagues used the sentence condition in the reading-based language localizer task (Fedorenko et al., 2010). Here, we attempted to extend this finding to auditory language processing and to a larger set of polyglots. To do so, we compared the responses in the language network to passages in one’s native language between our polyglot participants and a relatively large set of non-polyglot bilingual participants (n=86) who had previously completed a similar passage listening task in their native language (reported in Malik-Moraleda, Ayyash, et al. 2022; data available at https://osf.io/b9c4z). We fitted a model that predicted the BOLD response in the language network to the native language condition (relative to fixation), with Group (polyglot, control) modeled as a fixed effect and participants and fROIs modeled as random intercepts:

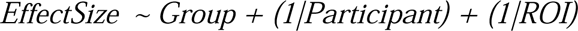

## Results

### 1. The polyglots’ language network responds to both familiar languages (of varying proficiency) and unfamiliar languages

As can be seen in **Figure 1**, each of the four familiar languages (L1 (Native language), L2, L3, and L4) elicited a response in the (individually defined) language areas that was stronger than the response to the quilted control condition (*p*s<0.001) (see **Figure S1** for L1-L4 activation overlap in all 34 participants and **Figure S2** for individual-language activation maps). Furthermore, unfamiliar related languages and unfamiliar unrelated languages also elicited a reliable response relative to the quilted control (*p*s<0.001). This response profile was present in each fROI individually (**Figure S3a, Table S4**) and it was similar when the language fROIs were defined using the L1 > Quilted contrast from the critical task (**Figure S4**).

**Figure 1.** Responses to different languages in the polyglots’ language network. **a.** Activation overlap for the four familiar languages (L1-L4). For each language, we selected 10% of voxels in the left hemisphere that were most responsive to the Language > Quilted-control contrast (based on the contrast values). The activations are shown within the boundaries of the language parcels (see <u>Methods</u>). Colors correspond to the *number of languages* (between 1 and 4) for which the voxel was in the set of top 10% of most responsive voxels. (Three-digit numbers in black boxes correspond to the unique ID of the participant and can be cross-referenced with the data on OSF: https://osf.io/3he75/.) For similar overlap maps for all participants, see **Figure S1**; for the activation maps for individual languages within sample participants, see **Figure S2**. **b.** Response in the language network (averaged across the five fROIs) to the conditions of the multi-language listening experiment relative to the fixation baseline (for the responses of the individual fROIs, see **Figure S3a**; for the responses broken down by experiment version (Bible, Alice in Wonderland), see **Figures S3b, S5c**). The language fROIs are defined by the Sentences > Nonwords contrast in the English localizer (see <u>Methods</u>; see **Figure S4** for evidence that the results are similar when the L1 > Quilted contrast from the critical task is used as the localizer). The conditions include the participant’s native language (L1), three languages that the participant is somewhat proficient in (L2, L3, and L4; proficiency is highest for L2, lower for L3, and lower for L4, as described in <u>Methods</u>), four unfamiliar languages (two languages that are related to the languages that the participant is relatively proficient in, two languages that the participant is completely unfamiliar with), and the perceptually matched control condition (Quilts; see <u>Methods</u>). Dots correspond to individual participants, error bars represent standard errors of the mean by participant.

### 2. Aside from their native language, the polyglots’ language network responds more strongly to higher-proficiency languages, and to unfamiliar languages that are related to high-to-moderate-proficiency languages than to unfamiliar unrelated languages

The response magnitudes varied across languages (**Figure 1b**), and this across-condition pattern was robust across participants and scanning runs (**Figure S5a-b**) and, largely, across the two sets of experimental materials (Bible stories vs. Alice in Wonderland passages; **Figure S5c**). The pattern also held when excluding the n=3 left-handed participants (**Figure S14**) and when using the RH (instead of the LH) language fROIs for the right-lateralized participant (**Figure S15**).

First, the response to the three familiar non-native languages (L2-L4) showed a gradient, with a stronger response to languages for which the polyglot reported higher proficiency. The response to L2 was numerically higher than the response to L3 (2.06 vs. 1.83 % BOLD signal change relative to fixation (here and elsewhere); β=0.236, n.s.), and the response to L3 was higher than the response to L4 (1.83 vs. 1.54; β=0.288, p=0.025). (This pattern held for the two sets of experimental materials (**Figure S5c**) and when excluding the participants (n=3) who were early bilinguals in their L1 and L2 (**Figure S7**).)

Second, the native language (L1) condition elicited a response that was reliably lower than the response to L2 (1.58 vs. 2.06; β=-0.483, p<0.001), in spite of the fact that L1 and L2 were rated similarly high in terms of proficiency (see <u>Methods</u>). (This pattern held when excluding the participants (n=3) for whom native language proficiency was below the maximum score of 20 (**Figure S6**).) However, the L2>L1 effect was not robust across the two sets of experimental materials: it appears to be driven by the subset of participants who listened to the Bible-stories materials (**Figure S5c**). For those participants, the effect is reliable (1.46 vs. 0.939; β=0.524, p<0.001) but for the participants who performed the Alice in Wonderland version of the experiment, the response to L2 is similar in magnitude to L1 (1.65 vs. 1.73; β=-0.081, n.s.). This difference between the two versions of the experiment may be due to the fact that the Bible-stories materials are linguistically simpler (use higher frequency words and constructions) than the Alice in Wonderland materials, based on the authors’ informal assessment of the materials for several of the languages. However, this difference was not predicted *a priori*, and a more complete understanding of this effect would require a within-participant evaluation of brain responses to materials that *only* differ in linguistic complexity and are more carefully matched across languages. Interestingly, in the participants who listened to the Bible-stories materials, the L2>L1 effect is more pronounced in the frontal compared to the temporal fROIs (**Figure S3b**), as evidenced by a condition (L1, L2) by ROI group (frontal, temporal) interaction (β=-0.770, p<0.05; main effect of condition: β=1.23, p<0.001; main effect of ROI group: β=0.997, p<0.001). Thus, for some polyglots, native language processing (at least for linguistically simple materials) can be carried out more focally, within the temporal lobe, with only minimal contribution from the frontal areas (cf. **Figures 2b, S10** for evidence of strong frontal responses during native language processing in non-polyglots).

**Figure 2.** Response in the language network to the native language in polyglots (purple, n=34) and non-polyglot bilinguals (grey, n=86). **a.** Response in the language network (averaged across the five fROIs). The brain inset shows the five parcels that were used for constraining the individual language fROI definition (see <u>Methods</u>). For the results in the subset of participants who were not included in Jouravlev et al. (2021), see **Figure S11**. **b.** Response in the five language fROIs separately. Dots correspond to individual participants, error bars represent standard errors of the mean by participant.

And third, unfamiliar related languages elicited a lower response than familiar languages (1.19 vs. 1.81; β=0.614, p<0.001), in line with the proficiency result observed for the familiar languages (L2>L3>L4). However, in spite of no past experience with *any* of the unfamiliar languages, unfamiliar related languages elicited a stronger response than unfamiliar unrelated languages (1.19 vs. 0.714; β=-0.481, p<0.001; this pattern also held when excluding participants (n=2) with errors in the selection of unfamiliar related languages (**Figure S8**; see **Table S6** caption for details)).

The pattern of response to the different languages was generally similar in the RH homotopic language network (**Figure S9**), but the magnitudes did not differentiate familiar and unfamiliar languages as clearly due to a relatively weaker response to the familiar languages in the RH compared to the LH language network.

### 3. Replication of Jouravlev et al. (2021): the polyglots’ language network responds less strongly during native language processing than the language network of non-polyglots

As can be seen in **Figure 2** (see also **Figure S10** for sample activation maps of polyglots and non-polyglot controls), we successfully replicate Jouravlev et al.’s (2021) finding in the auditory modality: polyglots showed a lower response in their language network while listening to their native language than a control group of 86 non-polyglot bilinguals (1.56 vs. 2.46 % BOLD signal change relative to the fixation baseline; β=-0.892, p<0.001). This effect held in all five language fROIs (p<0.01, uncorrected, **Table S5**) and numerically, for the subset of polyglots (n=19) that were not included in Jouravlev et al. (2021) (1.92 vs. 2.46; β=-0.539, p=0.06; see **Figure S11**). Note that all three participants who rated their native language proficiency lower than the maximum of 20 were excluded in these analyses.

## Discussion

The vast majority of humans grow up speaking or signing one or two languages. However, a small fraction of the population master a large number of languages (sometimes, several dozen). Although this phenomenon of polyglotism is not new (e.g., Erard, 2012), very few past studies have attempted to characterize the minds and brains of such individuals (e.g., Papagno & Vallar, 1995; Paradis, 2001; Amunts et al., 2004; Hervais-Adelman et al., 2015, 2018). The only prior functional brain imaging study that has investigated the language system of polyglots (Jouravlev et al., 2021) focused on comparing polyglots and non-polyglots during native language processing. Building on that work, we here took a deeper look at the polyglots’ language system and examined neural responses to several familiar and unfamiliar languages. We found that i) all languages, both familiar and unfamiliar, elicit a reliable response across the language network relative to a perceptually-matched control condition; ii) languages for which participants report higher proficiency levels elicit stronger responses; iii) the native language elicits a similar or lower response than a non-native language of similar proficiency; and iv) unfamiliar languages that are related to high-to-moderate proficiency languages elicit a reliably greater response than unfamiliar unrelated languages. In line with recent emphasis in the field on robustness and replicability (e.g., Ioannidis, 2005; Button et al., 2013; Ioannidis et al., 2014; Simmons et al., 2015; Poldrack et al., 2017), we also replicate Jouravlev et al.’s (2021) finding of lower responses during native language processing in polyglots compared to non-polyglots, extending the results to auditory language comprehension (cf. reading comprehension in the original study). Below, we elaborate on the significance of these findings and situate them within the broader empirical and theoretical landscape.

### All languages of a polyglot, and even unfamiliar languages, activate the language network

A set of interconnected frontal and temporal brain areas in the left hemisphere have long been known to support language processing. This “language network” supports the processing of typologically diverse languages (Malik-Moraleda, Ayyash et al., 2022; see e.g., Illes et al., 1999; Chee et al., 1999; Hernandez et al., 2001; Briellmann et al., 2004 for earlier evidence from smaller sets of languages), including even constructed languages, like Esperanto and Klingon (Malik-Moraleda et al., 2023). On the other hand, diverse non-linguistic inputs and tasks elicit little or no response in these brain areas (e.g., Fedorenko et al., 2011; Monti et al., 2012; Fedorenko et al., 2012; Deen et al., 2015; Pritchett et al., 2018; Jouravlev et al., 2019; Amalric & Dehaene, 2019; Ivanova et al., 2020; Liu et al., 2020; Chen et al., 2023; for reviews, see Fedorenko & Blank, 2020; Fedorenko et al., in press). Cross-linguistic universality and selectivity for language jointly suggest that some properties of the language network make it well-suited to support the broadly common features of languages that distinguish language processing from other cognitive processes. Perhaps not surprisingly then, when an individual acquires multiple languages (multiple sets of mappings between linguistic forms and meanings), those languages are represented and processed by the same neural system. This claim has been previously made based on fMRI data from bilingual and trilingual individuals (e.g., Illes et al., 1999a; Chee et al., 2000; Hernandez et al., 2001; Klein et al., 2006; Emmorey & McCullough, 2009; Buchweitz et al., 2009; Videsott et al., 2010; Willms et al., 2011; Honey et al., 2012; see Sebastian et al., 2011 and Sulpizio et al., 2020 for meta-analyses; see Perani & Abutalebi, 2005 and van Heuven & Dijkstra, 2010 for reviews). However, many past studies have used a) group-averaging analyses (cf. Dehaene et al., 1997), which may overestimate overlap, and/or b) paradigms that conflate linguistic and general task demands, which makes the nature of the overlap difficult to interpret. We used individual-subject fMRI analyses and an extensively validated language “localizer” paradigm, which effectively isolates language processing from task demands, to investigate neural responses to familiar and unfamiliar languages in a set of polyglots. We found that all languages that we examined (the native language, non-native languages of varying proficiency, and even unfamiliar languages) elicit a reliable response across the language network relative to a perceptually-matched control condition. Importantly, although our current results align with past research that reported inter-language overlap in bilingual individuals based on group-averaging analyses (e.g., Illes et al., 1999; Roux & Trétmoulet, 2002; Klein et al., 2006; Sulpizio et al., 2020), many examples exist in the literature where individual-subject analyses have overturned claims based on group-analytic approaches (e.g., see Fedorenko, 2021 for discussion); as a result, without conducting studies that take inter-individual topographic variability into account, one cannot be confident in the claims of activation overlap that come from group-averaging studies or meta-analyses of such studies. Moreover, by using a paradigm that robustly isolates language processing from task demands (e.g., Fedorenko & Blank, 2020), we can be confident that the overlap we observe has to do with linguistic processing specifically.

It is also important to note that the research questions that we focused on in the current study concern the core, left-hemisphere language network. As a result, the claim we are making here is a *positive* claim about all languages eliciting a response above the control condition in (all areas of) this core language network. We are not arguing that *no differences exist* among the languages in their activation topographies (for prior claims of such differences, see e.g., Dehaene et al., 1997; Fabbro et al., 2001; Xu et al., 2017; Correia et al., 2014). Such a claim would require a whole-brain analysis to establish that all languages engage all brain areas in a similar way (e.g., for any given area, either all languages elicit a reliable response, or none of the languages elicit a reliable response). We prefer to focus on the research questions that this study was designed to address and for which we have sufficient power, by virtue of being able to functionally localize our areas of interest (Nieto-Castanon & Fedorenko, 2012). However, we make all the individual whole-brain activation maps available (https://osf.io/3he75/), so other researchers can explore whole-brain topographic similarities/differences among the languages.

With respect to the core language network, it appears that in polyglots, processing *any* linguistic input—even if little/no meaning can be extracted from it—engages the language-processing mechanisms (see Malik-Moraleda, Ayyash et al., 2022 for evidence that this pattern also holds in non-polyglot bilingual individuals, with an unfamiliar foreign language condition eliciting a reliable response in the language network, albeit substantially weaker than the response to one’s native language). Of course, at a finer spatial scale, the different languages of a bilingual or multilingual individual must be dissociable (otherwise, there would be too much interference among the languages, making comprehension and production impossible). Indeed, a number of fMRI studies have reported reliable decoding of language identity in multivariate patterns of neural activity (Correia et al., 2014; Xu et al., 2017; Van de Putte et al., 2017; see Xu et al., 2021 for a review).

### What explains differences in the magnitude of the language network’s response to different languages?

The language network’s responses to the non-native languages—L2-L4 and the four unfamiliar languages—generally scale with self-reported proficiency. In particular, we observe a stronger response to familiar than unfamiliar languages (L2-L4 (moderate to high proficiency) > unfamiliar (no proficiency)), and a stronger response to familiar languages for which participants report higher proficiency levels (L2 > L3 > L4). However, two data points deviate from this proficiency-based scaling: i) the native language (L1), which has similar proficiency to L2 in this population (see <u>Methods</u>), elicits a lower response than L2 (although this effect is driven by the subset of the participants who performed the Bible-stories version of the experiment; see <u>Results</u>), and ii) among the unfamiliar languages, which would all receive a zero proficiency rating, languages that are related to high-to-moderate proficiency languages elicit a higher response than unfamiliar unrelated languages. What explains this overall pattern of results? There are at least two possibilities, which we discuss in turn.

#### Account 1

*The language network’s response to a particular language is a function of how strongly L1 (the native language) is activated*.

The first account is narrower in scope and originates in past work on bilingual language processing. In particular, a number of studies have suggested that native (L1) language representations are inevitably activated during the processing of non-native languages (e.g., L2 or L3) (e.g., Colomé, 2001; Thierry & Wu, 2007; Martin et al., 2009; Spalek et al., 2014; Giezen et al., 2015; Chen et al., 2017; Shook et al., 2019; Oppenheim et al., 2018; Jouravlev et al., 2020). With an additional stipulation that an asymmetry exists such that L2, L3, etc. are not/less co-activated during L1 processing (e.g., Costa & Santesteban, 2004), this account can explain the L2 > L1 effect (note that although this effect is not robust across the two sets of experimental materials in the current study (**Figure S5c**), it has been previously reported in the bilingual literature (e.g., de Bruin et al., 2014)). More specifically, the L2 > L1 effect could be due either i) to an overall greater number of linguistic representations being activated during L2 (or any other non-native language) processing (i.e., representations for both L1 and the input language; cf. only L1 representations activated during L1 processing), or ii) to the cost associated with suppressing L1 representations during non-native language processing. These two possibilities are difficult to differentiate empirically, as they make identical predictions in all cases as far as we can tell.

To explain the *L2 > L3 > L3 > unfamiliar* effect, this account would need to further stipulate that L1 is activated *more strongly* when an individual is processing higher-proficiency languages compared to lower-proficiency languages (e.g., Gollan, Slattery, Goldenberg et al., 2011). This is not an unreasonable stipulation: after all, individuals should typically have more experience with higher-proficiency languages and thus more opportunities to establish representational links with the corresponding words and constructions in L1. Alternatively, if L1 is activated to a similar extent during the processing of any non-native language (regardless of proficiency), then this account would need to include some additional factor that would explain the scaling of response with proficiency. It is also unclear how this account would explain the difference between the unfamiliar related and unfamiliar unrelated languages.

#### Account 2

*The language network’s response to a particular language is a function of a) the degree to which that language allows for the engagement of linguistic computations, and b) how costly that language is to process*.

The second account is broader in scope and finds support in studies outside of research on bilingualism/multilingualism. The general idea is that the level of the language network’s response to a stimulus can be explained with two factors: i) the degree to which the stimulus obeys the statistics of natural language, or the *language-like-ness* of the stimulus (responses are stronger to more language-like stimuli), and ii) *processing difficulty* (responses are stronger to linguistic input that is more difficult to process).

The first factor fits with the idea of ‘proper domains’ of specialized information processing systems (e.g., Sperber, 1994), whereby a stimulus needs to have certain properties—thus allowing for the engagement of particular computations—in order to engage the system. The language areas specialize for understanding and producing meaningful linguistic signals (e.g., Fedorenko et al., in press). For example, language comprehension requires computations related to phonological processing (recognizing familiar sounds and sound sequences), lexical processing (recognizing familiar word-forms and accessing their meanings), syntactic structure building (establishing dependencies among words using one’s grammatical knowledge), and semantic composition (deriving phrase- and sentence-level meanings). So, a stimulus whose processing requires these computations should elicit a high response, and a stimulus whose processing does not engage these computations should elicit a low response. In line with this prediction, past studies have found that, within a given language, a) sentences, which engage all of these computations, elicit a high response, b) linguistically degraded stimuli (like lists of unconnected words, ‘Jabberwocky’ sentences, or lists of pseudowords), which only engage some of these computations, elicit a lower response (e.g., Humphries et al., 2006; Fedorenko et al., 2010; Pallier et al., 2011; Bedny et al., 2011; Fedorenko et al., 2016; Shain, Kean et al., 2023), and c) non-linguistic inputs fail to engage the language network (e.g., Fedorenko et al., 2011; Monti et al., 2012; Deen et al., 2015; Pritchett et al., 2018; Jouravlev et al., 2019; Amalric & Dehaene, 2019; Ivanova et al., 2020; Liu et al., 2020, Chen et al., 2023; for reviews, see Fedorenko & Blank, 2020 and Fedorenko et al., in press).

The second factor is *processing difficulty*: when a stimulus falls within the proper domain of a system, responses are modulated by how difficult it is to process. For example, for language stimuli, the level of response in the language network can be predicted by both empirical measurements of incremental behavioral processing cost (e.g., word-by-word reading times; Henderson et al., 2020; Wehbe et al., 2021) and theoretical accounts of linguistic complexity, including surprisal-based accounts (e.g., Lopopolo et al., 2017; Shain, Blank et al., 2020; Heilbron et al., 2022) and memory-based accounts (e.g., Fiebach et al., 2001, 2002; Ben-Shachar et al., 2003; Santi & Grodzinsky, 2007; Blank et al., 2016; Shain et al., 2022).

In a recent study, Tuckute et al. (2024) recorded fMRI responses to 2,000 diverse linguistic strings and showed the influence of both these two factors: responses in the language network followed an inverted u-shape curve, with relatively low responses to non-language-like stimuli (resembling lists of unconnected words) and easy-to-process language stimuli (sentences that use common words and structures) and high responses to sentences that pose some processing cost due to having unusual structure and/or meaning.

Let us now apply this general framework to the current study. The processing of a language that an individual is highly proficient in (e.g., L2) should engage to the full extent all the computations that the language network supports, which should lead to a high response. A lower-proficiency language (e.g., L3 or L4) should engage a subset of these computations (e.g., phonological and lexical processes, with occasional or partial syntactic processing and semantic composition), which should therefore lead to a lower response. An unfamiliar language should engage phonological processes, and—if the language is related to one of the high-to-moderate proficiency languages—occasional lexical processes, which should lead to a low response. Finally, the native language (L1), similar to L2, should engage all the computations the network supports but may be relatively easier to process than L2, similar to simple, easy-to-process sentences within a language.

Although both accounts can explain some aspects of the data in the current study, the second account a) can explain the data without the need for various additional assumptions and stipulations, and b) is generally broader in scope. In particular, this account explains not only the level of response to different languages in polyglots, but also responses to linguistic and non-linguistic stimuli across many prior studies, as discussed above. This account can also explain lower responses to language in children relative to adults (e.g., Hiersche et al., 2023; Ozernov-Palchik, O’Brien et al., in prep.), and in individuals with stroke aphasia relative to age-matched healthy controls (e.g., Wilson & Schneck, 2020; Billot, 2023; Billot et al., 2023). Children are still gaining linguistic competence, actively learning new words and constructions, and individuals with aphasia have partially lost their linguistic knowledge or have difficulty accessing linguistic representations. As a result, during both language development and language loss, a linguistic stimulus may not fully engage the computations that it engages in a mature intact adult system, similar to how a low-proficiency language may not fully engage linguistic computations in polyglots. Given its explanatory breadth, the second account—whereby the level of response to a language is a function of how fully it can engage linguistic computations—is preferred to the account in terms of co-activation of the native language during the processing of non-native languages.

### The native language may hold a privileged status, at least in polyglots

The native language elicited a relatively low response in the language network. Although this finding was driven by the subset of the participants who performed the Bible-stories version of the experiment (see <u>Results</u>), it aligns with a) the finding reported in Jouravlev et al. (2021)—and replicated here—of lower responses to native language in polyglots compared to non-polyglots, and b) with several bodies of work that have argued that one’s native language may be represented and processed in a distinct way from the later acquired languages. For example, a) one’s native language has been shown to have unique and lasting advantages for processing speech in noisy conditions (e.g., Cutler, 2012; Blanco-Elorrieta et al., 2020); b) certain words, like taboo words and swear words, elicit stronger neural responses when presented in one’s native language (e.g., Chen et al., 2015; Sulpizio et al., 2019); and, more generally, c) linguistic content presented in one’s native language is processed in more emotional / less rational ways, as has been shown in the domains of economic and moral decision making (e.g., Keysar et al., 2012; Costa et al., 2014; Hayakawa et al., 2017).

Whether various previously reported effects of differential language processing in one’s native vs. non-native language relate to the lower magnitude of response to the native language in the left-hemisphere language network, or whether they instead, or in addition, relate to some other aspect(s) of the neural infrastructure of native language processing, remains to be determined. For example, although not the focus of the current paper, native and non-native languages appear to elicit differential responses in the fronto-parietal Multiple Demand (MD) network, which has been linked to executive control and goal-directed behaviors, and shown to be sensitive to processing difficulty across domains (e.g., Duncan, 2010, 2013; Fedorenko et al., 2013; Assem et al., 2020a). In particular, non-native, but not native, language processing, elicits above-baseline responses in the MD network (**Figure S12**; see also Malik-Moraleda, Ayyash et al., 2022, for evidence of below-baseline responses to one’s native language in the MD network of non-polyglot bilinguals; see Wolna et al., 2024 for evidence of stronger MD network engagement during non-native, compared to native, language processing). It is possible that the engagement of this network during non-native language processing is what leads to more rational responses to linguistic information, as reported in some past studies (e.g., Keysar et al., 2012; Costa et al., 2014; Hayakawa et al., 2017). This hypothesis can be tested in the future by relating the level of response in the MD network to the degree of rationality demonstrated in the economic/moral scenarios, although it would be necessary to first establish that the strength of the response to one’s non-native language in the MD network, as well as the relevant behavioral responses, are reliable within individuals and sufficiently variable across individuals. (Note that the *reason* for the MD network’s engagement during non-native language processing is at present unclear. Its engagement plausibly indicates greater processing difficulty associated with non-native language processing, but it remains unknown whether the MD network i) helps with linguistic computations (i.e., computations related to accessing word meanings and building syntactic structures), ii) provides additional attentional resources, iii) helps suppress L1 representations, which are putatively co-activated during non-native processing, as discussed in the preceding section, or iv) helps in some other way (e.g., Oppenheim, Dell and Schwartz, 2010).)

We also observed interesting differences between the frontal and temporal language regions in their response to native language processing. As can be seen from **Figure S3b**, it appears that—for the Bible-stories subset of the participants—the response to the native language is especially low in the frontal language areas, where it is similar in magnitude to the response to unfamiliar unrelated languages (cf. the temporal language areas, where it is much closer in magnitude to L2 and L3). This regional difference suggests that the native language (at least, when the materials are low in linguistic complexity) may be processed more focally, within the temporal component of the language network, with minimal involvement from the frontal language areas. If frontal language areas indeed play a less critical role in the processing of the native language in polyglots, then interfering with the activity of the inferior frontal areas in these individuals (e.g., via TMS; e.g., Devlin & Watkins, 2007) should have less of an effect on language performance, and damage to the inferior frontal areas should not have strong consequences on language function (see e.g., Luria, 1970; Kertesz & McCabe, 1977; Wilson et al., 2022 for evidence from non-polyglot individuals that frontal brain damage is *generally* less consequential than posterior temporal damage for aphasia).

### Replication of Jouravlev et al.’s (2021) finding of lower responses to language in polyglots compared to non-polyglots

Jouravlev et al. (2021) examined the language network in a set of 17 native-English-speaking polyglot individuals (13 of whom were included in the current study) and reported weaker and less spatially extensive responses during native language processing compared to both a set of carefully matched (on age, gender, and IQ) control participants and a larger set of controls. This effect was spatially selective: polyglots and controls did not differ in the strength or extent of activation in the right-hemisphere homotope of the language network or in two other large-scale networks (the Multiple Demand network and the Default network; Duncan, 2010, 2013; Assem et al., 2020a; Buckner & DiNicola, 2019). Here, we replicated and extended Jouravlev and colleagues’ results. In particular, whereas Jouravlev et al. (2021) relied on a reading-based language localizer paradigm, we examined the magnitude of response to *auditory* language comprehension and found reliably weaker responses in a set of 31 polyglots (for this analysis, we excluded the three participants who rated their native language proficiency lower than the maximum of 20). Thus, it appears that this finding is robust (although we acknowledge that other differences between our polyglot group and the control group could be contributing to the effect observed in the current study).

Jouravlev and colleagues interpreted their results as reflecting greater processing efficiency in polyglots and speculated that the difference between polyglots and non-polyglots is experientially driven. In particular, drawing on findings in the domain of motor learning (e.g., Poldrack et al. 1998; Fletcher et al. 1999; Kelly & Garavan 2005; Bernardi et al. 2013), they hypothesized that language representation and processing may become more efficient as a result of acquiring multiple languages. We find some support for this possibility in the form of a relationship between the strength of response to native language and the number of languages that a polyglot self-reported some proficiency in, such that individuals with more languages show weaker responses in the language network (**Figure S13**), with the proviso that n=34 is still a small sample for an individual-differences investigation, so this finding should only be considered suggestive. However, the possibility that individuals who become polyglots represent and process language more efficiently from the earliest stages of language acquisition remains a viable alternative.

Distinguishing between these possibilities will require genetic investigations of polyglots and/or longitudinal investigations of individuals as they acquire new languages. Investigating individuals who grow up in multilingual societies and learn multiple languages from birth (cf. individuals in the current study, who mostly grew up in monolingual societies and chose to learn their multiple languages later in life) can also inform the interpretation of the lower response to the native language in polyglots. If early multilinguals also show a lower response to their dominant/primary language(s) compared to matched mono/bilinguals, that would suggest that the effect is simply due to knowledge of multiple languages and thus unlikely to be genetic. If such individuals do not show a lower response to their dominant/primary language(s) compared to mono/bilinguals, that would suggest that the effect may be restricted to individuals who seek learning multiple languages and *may* have a genetic basis.

### Limitations

One limitation of the current study is that proficiency was estimated by self-report. In another recent study (Malik-Moraleda et al., 2023), we found that self-reported proficiency scores correlate with objective proficiency measures (see also Shameem, 1998; Martin et al., 2012; Diamond et al., 2014). Nevertheless, having an objective measure of proficiency remains an important desideratum for research on bi- and multilingualism. One challenge is having a measure that is validated and normed across a wide range of languages. Advances in AI and Natural Language Processing (NLP), including in multilingual neural models (MLNMs; e.g., Devlin et al., 2018), may aid in the development of such assessments, but as of right now, no standardized tool exists that would work for e.g., all the languages (n=34) tested in the current study.

Another limitation is that we did not directly assess the comprehension level for the specific materials that were used in the study, and use overall proficiency scores as a proxy. One way to do this would be to conduct a post-scanning behavioral session where participants are played the same passages that they listened to in the scanner, and are asked to a) transcribe each passage, and b) translate it into English (a language in which all participants were proficient), or into their native language, to the best of their ability. The resulting transcriptions and translations could be evaluated using NLP tools. Future work would benefit from directly relating measures of comprehensibility to neural response magnitudes.

Yet another limitation of the current study is that in this population, proficiency is correlated with age of acquisition (at r=-0.553; **Table S1**). Both proficiency and age of acquisition are predictive of comprehension, as measured behaviorally (e.g., Hartshorne et al., 2018). However, we favor proficiency as the key factor that explains the results in the current study for two reasons. First, as discussed above, the account that we outlined— where higher-proficiency languages engage linguistic computations to a greater degree— explains not only the current data, but also a wide range of effects outside of the literature on bi-/multilingualism. In contrast, any account that focuses on age of acquisition is likely to be narrower at scope. And second, one relevant data point in the current study concerns the difference between L3 and L4: although in our participants, both languages are relatively late-acquired (average age of acquisition >17 years: *M*=17.3 for L3 and *M*=20.1 for L4), L3 elicited a substantially and reliably higher response than L4. The proficiency difference between these languages is quite large (L3: *M*=13.5, *SD*=3.34; L4: *M*=9.15, *SD*=3.29) and seems more plausible as an explanation than a small difference in age of acquisition, especially given the late age of acquisition for both. Nevertheless, future work should i) attempt to dissociate proficiency and age of acquisition (by examining responses to early-acquired but forgotten/low-proficiency languages and late-acquired but high-proficiency languages) and ii) investigate (or control for) other factors, such as language exposure/familiarity and amount and type of language use, which have been previously shown to affect responses in the language areas (e.g., Wartenburger et al., 2003; Perani et al., 2003; De Bleser et al., 2003; Rodriguez Fronells et al., 2005; Wang et al., 2023).

### Summary and conclusions

The current fMRI investigation constitutes the first attempt to characterize the responses to different languages in the language network of polyglots and hyperpolyglots. Using a robust individual-subject approach and identifying language areas using a validated language localizer paradigm (Fedorenko et al., 2010), we uncovered several clear patterns. The main result is the scaling of the language network’s response magnitude with proficiency, with stronger responses to higher-proficiency languages. We argue that higher-proficiency languages elicit a higher response in the language network because they allow for the full range of linguistic computations to be engaged. Our findings contribute to the general understanding of the human language system and its ability to process familiar and unfamiliar languages in polyglot individuals.

## Supporting information

Supplementary Information for Malik-Moraleda, Jouravlev et al. (2024)

## Contributions

Conceptualization: SMM, OJ, KM, IB, EF

Methodology: SMM, OJ, ZM, KM, IB, EF

Software: ZM, TC

Investigation: all authors Formal analysis: SMM, OJ

Validation: SMM

Data curation: SMM, OJ, MT

Visualization: SMM, OJ, ZM

Writing - original draft: SMM, OJ, EF

Writing - review and editing: MT, ZM, TC, KM, IB

Supervision and project administration: EF

## Acknowledgements

We would like to acknowledge the Athinoula A. Martinos Imaging Center at the McGovern Institute for Brain Research at MIT, including its support team (Steve Shannon and Atsushi Takahashi). We would also like to thank i) Nancy Kanwisher for supporting pilot investigations for this study (back in 2013-14), ii) Josef Affourtit, Zoya Fan, Matt Siegelman, and Sophia Zhang for help with collecting and organizing the language background questionnaire data, iii) Richard Futrell and Shawn Wen for help in creating the Bible stories materials, iv) Sam Norman-Haignere for help in creating the quilted control clips, v) Zuzanna Balewski for help with experimental scripts, vi) members of the Fedorenko and Gibson labs for help with fMRI data collection (especially Eghbal Hosseini, Hope Kean, Anna Ivanova, and Lia Washington), vii) Judith Thurman, Yvonne Stapp, Simon Calder, Christian Saunders, Patrick Cox, Jessica Contrera, Gretchen McCulloch, Natalia Mesa, Kim Mills, Melina von Kivvon, and Susan Fitzgerald for covering this work in podcasts and news outlets and thus helping us recruit more polyglots, viii) Michael Erard, Simon Fisher, and Narly Golestani for helpful discussions, and of course, ix) our participants for their time and enthusiasm.

