## Supplementary Information for Malik-Moraleda, Jouravlev et al. (2024) for "Functional characterization of the language network of polyglots and hyperpolyglots with precision fMRI"

**Table S1:** Demographic details and the linguistic background of the participants.

**Table S2:** Language lateralization indices of the participants.

**Table S3:** The languages that were used in the experiment.

**Table S4:** A statistical comparison of response magnitudes for each of the language conditions vs. the quilt control condition in each language fROI separately.

**Table S5:** A statistical comparison of response magnitudes for the native language condition in polyglots and non-polyglot control participants in each language fROI separately.

**Table S6:** Information on the unfamiliar related languages (URLs) used in the experiment.

**Text S1:** Robustness of the results to the materials used (Bible vs. Alice).

**Text S2:** Robustness of the results to participants' age.

**Figure S1:** Activation overlap for the familiar languages (L1-L4) in all participants (n=34).

**Figure S2:** Activation overlap and individual language activations for the familiar languages (L1-L4).

**Figure S3:** Responses to different languages in the polyglots' language network shown for each fROI separately, including broken up by the materials used (Bible vs. Alice).

**Figure S4:** Responses to different languages in the polyglots' language network (defined by the L1 > Quilted contrast).

**Figure S5:** Robustness of the results across scanning runs, participants, and the materials used (Bible vs. Alice).

**Figure S6:** Responses to different languages in the polyglots' language network when excluding participants whose native language proficiency is below the maximum score of 20.

**Figure S7:** Responses to different languages in the polyglots' language network when excluding participants whose L2 is also a native language.

**Figure S8:** Responses to different languages in the polyglots' language network when excluding participants with errors in the selection of related languages.

**Figure S9:** Responses to different languages in the polyglots' right hemisphere homotope of the language network.

**Figure S10:** Activation maps for polyglots and non-polyglot control participants for native language processing.

**Figure S11:** Response to the native language in the language network of polyglots (purple, n=19 participants who were not included in Jouravlev et al., 2021) and non-polyglot controls (grey, n=86).

**Figure S12:** Responses to different language conditions in the polyglots' language and Multiple Demand (MD) networks.

**Figure S13:** Response to the native language as a function of the number of languages spoken by the polyglot.

**Figure S14:** Responses to different languages in the polyglots' language network when excluding left-handed participants.

**Figure S15:** Responses to different languages in the polyglots' language network in the language-dominant hemisphere.

| *UID* | *Gender* | *Age* | *Hand* | *nLangs* | *Languages* |
|---|---|---|---|---|---|
| *206* | f | 30 | l | 8 | [L1] English (0,20), [L2] Japanese (18,16), [L3] Spanish (12,11), T (24,7), P (27,6), C (11,6), K (28,6), [L4] Mandarin (11,4) |
| *320* | f | 28 | r | 5 | [L1] English (0,20), [L2] Japanese (19,20), [L3] Spanish (14,20), P (0,20), [L4] French (14,12) |
| *380* | m | 43 | r | 12 | [L1] English (0,20), [L2] Mandarin (20,19), [L3] Russian (25,12), J (21,10), G (NA,10), F (NA,10), [L4] Spanish (39,9), V (38,8), E (32,6), T (28,4), T (28,4), T (NA,4) |
| *383* | f | 19 | r | 5 | [L1] Spanish (0,20), [L2] English (0,20), [L3] Italian (13,19), F (18,15), [L4] Mandarin (13,13) |
| *385* | f | 29 | r | 10 | [L1] English (0,20), [L2] Japanese (17,19), [L3] French (12,17), K (21,15), [L4] Mandarin (23,10), S (27,10), H (28,8), I (28,8), A (27,4), M (26,4) |
| *389* | m | 29 | r | 16 | [L1] English (0,20), [L2] German (16,18), [L3] Spanish (12,16), [L4] Mandarin (20,14), H (NA,12), S (18,8), C (24,8), R (20,6), A (21,6), A (23,5), J (28,5), N (27,5), T (27,4), M (23,4), K (NA,3), T (NA,3) |
| 394 | m | 24 | r | 5 | [L1] French (0,20), [L2] English (11,20), [L3] Russian (25,16), [L4] Spanish (13,15), P (21,NA) |
| 396 | m | 29 | r | 8 | [L1] French (0,20), [L2] Spanish (12,20), [L3] English (10,19), [L4] Italian (NA,7), C (NA,NA), S (NA,NA), P (NA,NA), N (NA,NA) |
| 439 | m | 32 | r | 7 | [L1] English (0,20), [L2] German (16,20), [L3] Arabic (21,18), [L4] Italian (25,16), H (29,13), P (26,8), F (3,6) |
| 452 | m | 29 | r | 54 | [L1] English (0,20), [L2] Mandarin (16,19), K (14,16), [L3] Japanese (10,13), V (18,13), [L4] Spanish (6,10), F (14,10), A (18,9), R (15,9), C (14,9), T (21,8), G (12,8), I (17,8), M (16,7), E (18,7), B (16,7), D (14,7), E (22,6), F (18,6), G (19,6), A (25,6), C (18,6), H (20,6), H (16,6), M (18,6), I (18,6), K (18,5), L (18,5), D (22,5), I (19,5), I (18,5), M (25,5), N (18,5), P (18,5), P (17,5), T (20,5), T (18,5), P (28,4), P (20,4), P (20,4), Q (18,4), S (18,4), X (25,4), Y (28,4), Z (25,4), C (28,4), T (25,4), W (18,4), B (18,4), H (18,4), N (18,4), S (18,4), L (18,4), B (NA,4) |

| | | | | | |
|---|---|---|---|---|---|
| 511 | f | 30 | a | 12 | [L2] English (9,20), [L3] French (12,14), [L1] Mandarin (0,12), [L4] Italian (28,11), S (NA,NA), P (NA,NA), A (NA,NA), A (NA,NA), L (NA,NA), G (NA,NA), A (NA,NA), Y (NA,NA) |
| 657 | f | 51 | r | 26 | [L1] English (0,20), [L2] Spanish (5,16), G (2,16), F (NA,16), [L3] Italian (44,12), P (NA,11), P (NA,11), G (NA,9), P (NA,8), [L4] Arabic (43,7), Urdu (NA,7), R (NA,7), D (NA,6), H (NA,6), N (NA,6), S (NA,5), J (NA,5), M (NA,5), Y (NA,5), K (NA,5), I (NA,5), P (NA,5), S (NA,4), L (NA,4), U (NA,4), T (NA,4) |
| 663 | m | 43 | r | 13 | [L1] English (0,20), [L2] Spanish (5,16), B (19,16), F (17,8), [L3] Romanian (42,6), R (18,6), C (0,5), D (23,5), G (18,5), [L4] Turkish (0,4), H (17,4), M (18,4), J (24,4) |
| 668 | m | 39 | r | 29 | [L1] English (0,20), [L2] Mandarin (17,19), [L3] Indonesian (17,16), P (NA,16), P (NA,16), M (NA,16), I (NA,16), S (NA,16), A (NA,14), W (NA,14), [L4] French (12,13), S (NA,13), B (NA,13), O (NA,12), A (NA,12), W (NA,12), S (NA,12), B (NA,10), M (NA,10), I (NA,9), B (NA,8), C (NA,8), F (NA,8), A (NA,8), B (NA,8), C (NA,8), M (NA,8), A (NA,8), Q (NA,6) |
| 701 | m | 59 | r | 13 | [L1] English (0,20), [L2] Mandarin (19,12), [L3] Japanese (23,8), [L4] German (30,8), P (NA,NA), F (11,NA), R (17,NA), M (25,NA), M (39,NA), S (57,NA), Italian (NA,NA), P (NA,NA), S (NA,NA) |
| 798 | f | 42 | r | 11 | [L1] English (0,20), [L2] Russian (0,20), [L3] Italian (19,8), [L4] Arabic (26,8), S (15,NA), P (21,NA), F (15,NA), I (23,NA), L (35,NA), S (NA,NA), H (NA,NA), G (NA,NA) |
| 834 | f | 28 | r | 7 | [L1] Spanish (0,20), [L2] English (6,20), C (6,20), K (0,15), F (12,14), [L3] Hindi (10,12), [L4] Arabic (25,8) |
| 839 | m | 31 | r | 7 | [L2] English (4,20), [L1] Hungarian (0,10), T (0,10), [L3] Spanish (14,10), [L4] French (6,4), A (6,4), R (17,4) |
| 867 | m | 46 | r | 35 | [L1] English (0,20), [L2] Spanish (0,20), R (12,20), C (14,20), S (15,20), B (16,18), R (14,18), P (14,18), S (16,18), N (22,18), F (15,17), N (20,17), P (15,17), U (16,16), C (20,16), I (27,15), L (17,15), I (20,15), G |

| | | | | | |
|---|---|---|---|---|---|
| | | | | | (11,15), E (19,13), S (14,12), G (15,12), M (25,12), [L3] Dutch (14,12), [L4] Arabic (15,12), S (40,12), A (14,12), W (44,12), F (7,12), G (26,12), N (40,11), Z (29,10), L (27,8), M (33,6), J (20,6) |
| 986 | f | 67 | r | 5 | [L1] English (0,20), [L2] Hebrew (15,17), F (7,11), [L3] Italian (2,9), [L4] German (19,6) |
| 990 | f | 57 | r | 48 | [L1] English (0,20), S (5,20), F (6,20), [L2] Russian (23,20), D (26,16), [L3] Portuguese (26,16), G (5,16), S (40,16), S (26,16), S (20,16), G (19,14), I (40,12), Y (56,12), [L4] Norwegian (20,12), A (43,12), H (26,12), U (56,12), L (13,10), D (19,10), B (24,10), C (56,8), Z (42,8), J (56,8), M (55,8), H (56,8), A (34,8), K (32,8), I (23,8), S (56,8), V (56,8), L (46,8), L (46,8), W (24,8), I (24,8), T (23,8), A (26,8), C (24,8), P (24,8), R (24,8), T (26,8), H (24,8), F (20,8), H (26,8), F (20,8), N (56,8), W (23,8), F (26,8), C (24,8) |
| 993 | m | 57 | l | 8 | [L1] English (0,20), [L2] French (13,18), S (15,17), [L3] German (50,12), P (57,11), R (33,10), I (40,9), [L4] Korean (56,8) |
| 994 | f | 22 | r | 11 | [L1] German (0,20), [L2] English (10,18), N (18,14), [L3] Dutch (20,13), [L4] Italian (19,9), F (9,7), M (17,6), D (20,6), Q (22,4), S (22,4), B (22,4) |
| 995 | m | 29 | r | 10 | [L1] Dutch (0,20), [L2] English (8,19), N (18,15), [L3] German (12,14), [L4] French (12,9), J (20,9), L (12,5), A (12,5), I (18,5), L (17,4) |
| 996 | m | 31 | r | 21 | [L1] English (0,20), S (20,20), [L2] Spanish (9,18), F (16,18), W (22,15), [L3] Irish (20,15), [L4] German (19,14), I (16,13), P (NA,10), J (13,9), D (NA,9), G (14,8), A (27,8), G (16,8), H (27,8), T (30,8), C (NA,6), L (12,6), K (18,6), S (18,6), M (NA,5) |
| 997 | m | 24 | r | 35 | [L1] English (0,20), [L2] Spanish (8,20), P (15,16), I (16,16), F (16,16), R (16,12), [L3] Russian (16,12), C (16,12), T (18,12), B (24,12), [L4] Dutch (22,10), G (19,8), P (0,8), U (22,8), C (20,8), M (18,8), L (18,6), G (22,6), B (22,6), H (18,6), U (18,6), P (20,6), S (18,6), L (18,5), N (22,5), D (22,5), S (22,5), B (18,5), C (20,5), H (18,4), A (18,4), A (18,4), T (18,4), J (18,4), K (6,4) |

| | | | | | |
|---|---|---|---|---|---|
| 1012 | m | 32 | r | 21 | [L1] English (0,20), [L2] Spanish (9,20), P (19,20), H (18,19), F (8,17), G (5,16), I (9,15), S (22,15), [L3] Japanese (15,12), F (30,10), B (32,9), [L4] Mandarin (18,8), Z (22,8), T (20,8), Y (20,8), P (32,7), A (22,7), T (20,6), B (20,5), I (25,4), Q (21,4) |
| 1022 | m | 24 | r | 7 | [L1] French (0,20), E (15,17), [L2] Spanish (11,16), [L3] English (13,14), I (15,11), P (15,10), [L4] Greek (20,6) |
| 1026 | m | 21 | r | 22 | [L1] English (0,20), [L2] Japanese (10,17), F (11,17), R (12,17), S (8,17), A (0,15), M (15,14), [L3] Korean (15,12), I (14,11), U (19,10), G (17,9), A (6,9), P (19,9), V (20,8), H (18,8), [L4] Portuguese (18,8), C (18,7), T (18,6), K (19,6), C (16,6), L (19,6), H (19,4) |
| 1032 | f | 26 | r | 6 | [L1] Russian (0,20), [L2] English (11,20), [L3] Italian (16,14), S (26,8), [L4] Hebrew (7,7), H (19,6) |
| 1033 | m | 27 | r | 7 | E (9,20), [L2] Hebrew (3,20), [L1] Russian (0,19), [L3] Arabic (12,11), S (9,10), [L4] French (9,9), F (15,8) |
| 1034 | f | 20 | r | 5 | [L1] Russian (0,20), [L2] English (3,20), F (5,17), [L3] Spanish (7,16), [L4] German (12,11) |
| 1035 | f | 20 | r | 5 | [L1] French (0,20), [L2] English (6,20), [L3] Spanish (10,16), K (0,7), [L4] Mandarin (11,5) |
| 1041 | m | 71 | r | 6 | [L1] English (0,20), [L2] Portuguese (26,19), P (26,16), [L3] Spanish (3,14), D (32,4), [L4] German (53,4) |

**Table S1: Demographic details and the linguistic background of the participants**. ***UID*** refers to the Unique ID assigned to the participant in the lab's database (and can be cross-referenced with the data available at OSF: https://osf.io/3he75/). ***Age*** refers to the participant's age (in full years) at the time of testing. ***Hand*** refers to the participant's handedness (r=right-handed; l=left-handed; a=ambidextrous). ***nLangs*** refers to the number of languages that the participant listed as having some proficiency in, and ***Languages*** lists all these languages. For each participant, the languages are ordered by (self-reported) proficiency. Familiar languages that were tested during the fMRI experiment (L1, L2, L3 and L4) are marked in red in square brackets before the relevant language (see Methods for details). In order to protect the participants' identities, for languages not tested in the experiment, we provide only the initial letter of the language (e.g., "E" for "English" or "Estonian"). The only exceptions (where we spelled out the languages) are cases where an unfamiliar related language was related to a language that was not tested in the experiment for that participant and is therefore listed in Table S6. Each language also has a parenthetical after it (e.g., "(x, y"): the first number corresponds to the *age of acquisition* and the second number corresponds to overall self-rated proficiency (as described in Methods, participants were asked to rate their ability in auditory comprehension, written comprehension, spoken production, and written production on a scale from 0 (no knowledge) to 5 (native or native-like proficiency); the scores were summed to derive an overall score (maximum value: 20)). (Note that age of acquisition and self-reported proficiency are correlated in this population, at r=-0.553.) Missing values are marked as NAs. A csv version of the table is available at OSF (https://osf.io/3he75/).

| UID | LH Suprathreshold Voxels | RH Suprathreshold Voxels | Lateralization Index |
|---|---|---|---|
| *206* | 1233 | 1233 | 0 |
| *320* | 1169 | 225 | 0.677 |
| *380* | 1011 | 404 | 0.429 |
| *383* | 819 | 379 | 0.367 |
| *385* | 1734 | 1566 | 0.051 |
| *389* | 755 | 138 | 0.691 |
| *394* | 2055 | 135 | 0.877 |
| *396* | 3432 | 1161 | 0.494 |
| *439* | 299 | 23 | 0.857 |
| *452* | 1616 | 988 | 0.241 |
| *511* | 117 | 1486 | -0.854 |
| *657* | 1999 | 625 | 0.524 |
| *663* | 560 | 808 | -0.181 |
| *668* | 1254 | 1290 | -0.014 |
| *701* | 2155 | 944 | 0.391 |
| *798* | 516 | 41 | 0.853 |
| *834* | 758 | 60 | 0.853 |
| *839* | 2971 | 1875 | 0.226 |
| *867* | 786 | 332 | 0.406 |
| *986* | 2206 | 1059 | 0.351 |
| *990* | 493 | 300 | 0.243 |
| *993* | 1297 | 471 | 0.467 |
| *994* | 1480 | 54 | 0.930 |
| *995* | 2263 | 441 | 0.674 |
| *996* | 2678 | 873 | 0.508 |
| *997* | 2594 | 2327 | 0.054 |
| *1012* | 1352 | 135 | 0.818 |
| *1022* | 2086 | 396 | 0.681 |
| *1026* | 1599 | 276 | 0.706 |
| *1032* | 1511 | 426 | 0.56 |
| *1033* | 1961 | 961 | 0.342 |
| *1034* | 3360 | 900 | 0.577 |
| *1035* | 2746 | 1288 | 0.361 |
| *1041* | 2144 | 868 | 0.424 |

**Table S2: Language lateralization indices of the participants**. ***UID*** refers to the Unique ID assigned to the participant in the lab's database (and can be cross-referenced with the data available at OSF: https://osf.io/3he75/). ***LH*** and ***RH Suprathreshold Voxels*** refer to the number of voxels within the boundaries of the language parcels (see Methods) summed across the five parcels in the left hemisphere (LH) and the right hemisphere (RH) that are significant for the language localizer contrast at a fixed ($p<0.001$ uncorrected whole-brain) statistical threshold. To calculate the ***Lateralization Index***, we used the following formula “LI = (number of LH voxels–- number of RH voxels) / (number of LH voxels + number of RH voxels)”.

| | Native | Familiar | | | Unfamiliar Related | | Unfamiliar unrelated | |
|---|---|---|---|---|---|---|---|---|
| **UID** | **L1** | **L2** | **L3** | **L4** | **L5** | **L6** | **L7** | **L8** |
| **206** | English | Japanese | Spanish | Mandarin | German | Italian | Basque | Georgian |
| **320** | English | Japanese | Spanish | French | Italian | German | Basque | Georgian |
| **380** | English | Mandarin | Russian | Spanish | Dutch | Italian | Basque | Georgian |
| **383** | Spanish | English | Italian | Mandarin | Italian | German | Basque | Georgian |
| **385** | English | Japanese | French | Mandarin | Italian | German | Basque | Georgian |
| **389** | English | German | Spanish | Mandarin | Italian | Dutch | Basque | Georgian |
| **394** | French | English | Russian | Spanish | Italian | German | Basque | Georgian |
| **396** | French | Spanish | English | Italian | Romanian | German | Basque | Georgian |
| **439** | English | German | Arabic | Italian | Dutch | Romanian | Basque | Georgian |
| **452** | English | Mandarin | Japanese | Spanish | Romanian | Yiddish | Basque | Georgian |
| **511** | Mandarin | English | French | Italian | Dutch | Romanian | Basque | Georgian |
| **657** | English | Spanish | Italian | Arabic | Romanian | Bangla | Basque | Georgian |
| **663** | English | Spanish | Romanian | Turkish | Italian | Yiddish | Basque | Georgian |
| **668** | English | Mandarin | Indonesian | French | German | Romanian | Thai | Georgian |
| **701** | English | Mandarin | Japanese | German | Dutch | Romanian | Basque | Georgian |
| **798** | English | Russian | Italian | Arabic | Dutch | Romanian | Basque | Georgian |
| **834** | Spanish | English | Hindi | Arabic | Italian | Dutch | Georgian | Thai |
| **839** | Hungarian | English | Spanish | French | Dutch | Norwegian | Tamil | Basque |
| **867** | English | Spanish | Dutch | Arabic | Georgian | Mandarin | Basque | Thai |
| **986** | English | Hebrew | Italian | German | Dutch | Arabic | Tamil | Basque |
| **990** | English | Russian | Portuguese | Norwegian | Ukrainian | Afrikaans | Tagalog | Tamil |
| **993** | English | French | German | Korean | Dutch | Catalan | Tamil | Basque |
| **994** | German | English | Dutch | Italian | Norwegian | Afrikaans | Tamil | Basque |

| | | | | | | | | |
|---|---|---|---|---|---|---|---|---|
| **995** | Dutch | English | German | French | Swedish | Afrikaans | Tamil | Basque |
| **996** | English | Spanish | Irish | German | Norwegian | Afrikaans | Tamil | Basque |
| **997** | English | Spanish | Russian | Dutch | Afrikaans | Belarusian | Tamil | Tagalog |
| **1012** | English | Spanish | Japanese | Mandarin | Dutch | Norwegian | Belarusian | Basque |
| **1022** | French | Spanish | English | Greek | Romanian | Dutch | Tamil | Basque |
| **1026** | English | Japanese | Korean | Portuguese | Dutch | Swedish | Tagalog | Basque |
| **1032** | Russian | English | Italian | Hebrew | Ukrainian | Dutch | Tamil | Basque |
| **1033** | Russian | Hebrew | Arabic | French | Ukrainian | Polish | Tamil | Basque |
| **1034** | Russian | English | Spanish | German | Dutch | Ukrainian | Tamil | Basque |
| **1035** | French | English | Spanish | Mandarin | Italian | Romanian | Tamil | Basque |
| **1041** | English | Portuguese | Spanish | German | Dutch | NA | Basque | NA |

**Table S3: The languages that were used in the experiment.** ***UID*** refers to the Unique ID assigned to the participant in the lab's database (and can be cross-referenced with the data available at OSF: https://osf.io/3he75/). For the familiar languages, L2 was the non-native language that participants reported being most proficient in; L3 and L4 were selected so that participants were somewhat proficient in them (a score of at least 4 out of 20 in each), but these were not always the next most proficient languages due to the limitations on the languages for which experimental materials were available. The unfamiliar related languages were selected to be related to L1 and L2 when possible. If not possible (in cases when the participant had familiarity with the available related languages or when the materials were not available), languages related to L3 or L4 were used (and in two cases, we used languages that are related to languages that were not tested as L1, L2, L3 or L4 but that the participant had proficiency in). (For two additional participants (UIDs 383 and 867), there were errors in the selection of the related languages, as elaborated in the caption for Figure S8. Finally, participant UID 1041 was tested for a different study and, as a result, was scanned on L1-L4, and only one unfamiliar related and one unfamiliar unrelated language; instead, the set of languages for this participant included two additional familiar languages—conditions that we excluded in order to adhere to the design of the experiment used for all other participants.)

| fROI | Condition Name | Critical condition Effect Size | Control condition (Quilts) Effect Size | Beta Value | P-Value |
|---|---|---|---|---|---|
| *LH IFGorb* | L1 | 1.06 | 0.012 | 1.05 | **<0.001** |
| *LH IFGorb* | L2 | 1.68 | 0.012 | 1.67 | **<0.001** |
| *LH IFGorb* | L3 | 1.32 | 0.012 | 1.30 | **<0.001** |
| *LH IFGorb* | L4 | 1.10 | 0.012 | 1.09 | **0.004** |
| *LH IFGorb* | L5 | 0.756 | 0.012 | 0.744 | 0.012 |
| *LH IFGorb* | L6 | 0.666 | 0.012 | 0.651 | **0.001** |
| *LH IFGorb* | L7 | 0.176 | 0.012 | 0.164 | 0.518 |
| *LH IFGorb* | L8 | 0.431 | 0.012 | 0.416 | 0.105 |
| *LH IFG* | L1 | 1.59 | 0.271 | 1.32 | **<0.001** |
| *LH IFG* | L2 | 2.25 | 0.271 | 1.98 | **<0.001** |
| *LH IFG* | L3 | 1.95 | 0.271 | 1.68 | **<0.001** |
| *LH IFG* | L4 | 1.92 | 0.271 | 1.65 | **<0.001** |
| *LH IFG* | L5 | 1.64 | 0.271 | 1.37 | **<0.001** |
| *LH IFG* | L6 | 1.45 | 0.271 | 1.15 | **<0.001** |
| *LH IFG* | L7 | 0.842 | 0.271 | 0.572 | 0.049 |
| *LH IFG* | L8 | 1.08 | 0.271 | 0.788 | **0.008** |
| *LH MFG* | L1 | 1.17 | -0.056 | 1.23 | **<0.001** |
| *LH MFG* | L2 | 1.76 | -0.056 | 1.82 | **<0.001** |
| *LH MFG* | L3 | 1.92 | -0.056 | 1.98 | **<0.001** |
| *LH MFG* | L4 | 1.57 | -0.056 | 1.63 | **<0.001** |
| *LH MFG* | L5 | 1.23 | -0.056 | 1.28 | **<0.001** |
| *LH MFG* | L6 | 0.920 | -0.056 | 0.980 | **0.002** |
| *LH MFG* | L7 | 0.495 | -0.056 | 0.551 | 0.075 |
| *LH MFG* | L8 | 0.716 | -0.056 | 0.776 | 0.013 |
| *LH AntTemp* | L1 | 1.94 | 0.369 | 1.57 | **<0.001** |
| *LH AntTemp* | L2 | 2.17 | 0.369 | 1.80 | **<0.001** |
| *LH AntTemp* | L3 | 1.80 | 0.369 | 1.43 | **<0.001** |
| *LH AntTemp* | L4 | 1.39 | 0.369 | 1.02 | **<0.001** |
| *LH AntTemp* | L5 | 1.35 | 0.369 | 0.983 | **<0.001** |
| *LH AntTemp* | L6 | 1.19 | 0.369 | 0.822 | **<0.001** |
| *LH AntTemp* | L7 | 0.737 | 0.369 | 0.367 | 0.035 |
| *LH AntTemp* | L8 | 0.907 | 0.369 | 0.545 | **0.002** |
| *LH PostTemp* | L1 | 2.13 | 0.400 | 1.73 | **<0.001** |
| *LH PostTemp* | L2 | 2.45 | 0.400 | 2.05 | **<0.001** |
| *LH PostTemp* | L3 | 2.14 | 0.400 | 1.74 | **<0.001** |
| *LH PostTemp* | L4 | 1.71 | 0.400 | 1.31 | **<0.001** |

| | | | | | |
|---|---|---|---|---|---|
| *LH PostTemp* | L5 | 1.39 | 0.400 | 0.988 | **<0.001** |
| *LH PostTemp* | L6 | 1.17 | 0.400 | 0.751 | **0.001** |
| *LH PostTemp* | L7 | 0.690 | 0.400 | 0.690 | 0.208 |
| *LH PostTemp* | L8 | 0.893 | 0.400 | 0.477 | 0.041 |

**Table S4: A statistical comparison of response magnitudes for each of the language conditions vs. the quilt control condition in each language fROI separately.** The following LME was fit for each ROI individually: *EffectSize ~ Condition + (1|Participant)*. The effect sizes for the critical and control conditions are relative to the fixation baseline. We report uncorrected *p*-values, but in **bold** we highlight the *p*-values that would survive the Bonferroni correction for five fROIs (these include—for all fROIs—L1 through L4, as well as L6; for all but the LH IFGorb fROI—L5; and for some fROIs—L7 or L8).

| fROI | Mean in polyglots | Mean in non-polyglot controls | Beta | P-value |
|---|---|---|---|---|
| *LH IFGorb* | 1.05 | 2.01 | -0.95 | **0.003** |
| *LH IFG* | 1.60 | 2.67 | -1.08 | **0.002** |
| *LH MFG* | 1.20 | 2.33 | -1.14 | **0.003** |
| *LH AntTemp* | 1.94 | 2.56 | -0.631 | **0.001** |
| *LH PostTemp* | 2.05 | 2.72 | -0.669 | 0.018 |

**Table S5: A statistical comparison of response magnitudes for the native language condition in polyglots and non-polyglot control participants in each language fROI separately.** We report uncorrected *p*-values, but in **bold** we highlight the *p*-values that would survive the Bonferroni correction for five fROIs (these include all but the LH PostTemp fROI).

| UID | Unfamiliar Related Language 1 (URL 1) | | | | Unfamiliar Related Language 2 (URL 2) | | | |
|---|---|---|---|---|---|---|---|---|
| | URL1 | Language to which URL1 is related (URL_Rel1) | URL_rel 1 actual language | Distance between URL1 and URL_Rel1 | URL2 | Language to which URL2 is related (URL_Rel2) | URL_rel 2 actual language | Distance between URL2 and URL_Rel2 |
| ***206*** | German | L1 | English | 31.3 | Italian | L3 | Spanish | 14.0 |
| ***320*** | German | L1 | English | 31.3 | Italian | L3 | Spanish | 14.0 |
| ***380*** | Dutch | L1 | English | 21.8 | Italian | L4 | Spanish | 14.0 |
| ***383*** | Italian (selected in error) | NA | NA | NA | German | L2 | English | 31.3 |
| ***385*** | German | L1 | English | 31.3 | Italian | L3 | French | 20.2 |
| ***389*** | Dutch | L1 | English | 21.8 | Italian | L3 | Spanish | 14.0 |
| ***394*** | Italian | L1 | French | 20.2 | German | L2 | English | 31.3 |
| ***396*** | Romanian | L1 | French | 37.7 | German | L3 | English | 31.3 |
| ***439*** | Dutch | L1 | English | 21.8 | Romanian | L4 | Italian | 25.0 |
| ***452*** | Yiddish | L1 | English | 38.3 | Romanian | L4 | Spanish | 33.0 |
| ***511*** | Dutch | L2 | English | 21.8 | Romanian | L4 | Italian | 25.0 |
| ***657*** | Romanian | L2 | Spanish | 33.0 | Bangla | NotTested (proficiency: 7) | Urdu | 21.9 |
| ***663*** | Yiddish | L1 | English | 38.3 | Italian | L2 | Spanish | 14.0 |
| ***668*** | German | L1 | English | 31.3 | Romanian | L4 | French | 37.7 |
| ***701*** | Dutch | L1 | English | 21.8 | Romanian | NotTested (proficiency: N/A) | Italian | 25.0 |
| ***798*** | Dutch | L1 | English | 21.8 | Romanian | L3 | Italian | 25.0 |
| ***834*** | Italian | L1 | Spanish | 14.0 | Dutch | L2 | English | 21.8 |
| ***839*** | Dutch | L2 | English | 21.8 | Norwegian | L2 | English | 28.3 |
| ***986*** | Dutch | L1 | English | 21.8 | Arabic | L2 | Hebrew | 27.9 |
| ***990*** | Afrikaans | L1 | English | 22.5 | Ukrainian | L2 | Russian | 8.4 |
| ***993*** | Dutch | L1 | English | 21.8 | Catalan | L2 | French | 20.5 |
| ***994*** | Afrikaans | L1 | German | 16.3 | Norwegian | L2 | English | 28.3 |
| ***995*** | Swedish | L1 | Dutch | 20.7 | Afrikaans | L2 | English | 22.5 |
| ***996*** | Norwegian | L1 | English | 28.3 | Afrikaans | L4 | German | 16.3 |
| ***997*** | Afrikaans | L1 | English | 22.5 | Belarusian | L3 | Russian | 10.6 |
| ***1012*** | Dutch | L1 | English | 21.8 | Norwegian | L1 | English | 28.3 |
| ***1022*** | Romanian | L1 | French | 37.7 | Dutch | L3 | English | 21.8 |

| | | | | | | | | |
|---|---|---|---|---|---|---|---|---|
| ***1026*** | Dutch | L1 | English | 21.8 | Swedish | L1 | English | 31.0 |
| ***1032*** | Ukrainian | L1 | Russian | 8.40 | Dutch | L2 | English | 21.8 |
| ***1033*** | Ukrainian | L1 | Russian | 8.40 | Polish | L1 | Russian | 7.60 |
| ***1034*** | Dutch | L2 | English | 21.8 | Ukrainian | L1 | Russian | 8.40 |
| ***1035*** | Italian | L1 | French | 20.2 | Romanian | L3 | Spanish | 33.0 |
| ***1041*** | Dutch | L1 | English | 21.8 | NA | NA | NA | NA |

**Table S6: Information on the unfamiliar related languages (URLs) used in the experiment.** For each of the two URLs (URL1 and URL2, both marked in red font), we provide information on i) the language condition to which the URL is related (e.g., L1, L2, etc.), ii) the actual language to which the URL is related (e.g., English), and iii) the distance between the URL and the language to which it is related, using the distance metric introduced in Beaufils & Tomin (2020), as detailed below.

***Participants that were affected by errors in the selection of URLs:***

→ Participant UID 383: For this participant, Italian was used as one of the URLs. However, this is one of their familiar languages and indeed a language that was used as condition L3.

→ Participant UID 867 (not included in this table): Given that this participant spoke too many languages, it was impossible to identify any URLs within the sets of languages that were available for testing. (We ended up using Georgian and Mandarin, which were both familiar to the participant. We excluded these data points from our analyses.)

→ Participant UID 1041: For this participant, only one URL was included (see the caption for Table S3).

***Details of the inter-language distance metric:*** Beaufils & Tomin (2020) developed a metric based on comparing lexical entries across >200 diverse languages (http://www.elinguistics.net/Compare_Languages.aspx). The scale goes from 1 to 100, with distance values 1-30 being considered highly related languages; 30-50 considered related languages (with a shared protolanguage between 2,000 and 4,000 years ago), between 50-70 considered remotely related languages (with a shared proto-language approximately between 400 and 6,000 years ago), between 70 and 78 considered very remotely related language, and between 79 and 100 considered unrelated.

To statistically compare the distances between a) unfamiliar related languages (URLs) and their associated languages (columns 3 and 7 in the table above; a total of n=64 data points—see above for exclusions), and b) unfamiliar unrelated languages (UULs; n=67 data points—see above for exclusions) and high-proficiency languages, we performed the following analysis:

First, for the UULs, we extracted the distance values between each language and L1, L2, L3, and L4 for each participant, and then averaged across these four values to obtain a single value per UUL per participant. Next, we compared the two sets of distance values (URLs (n=64): $M$=23.25; $SD$=7.95; and UULs (n=67): $M$=89.9, $SD$=2.99) using an independent-samples $t$-test, which revealed a highly reliable difference ($t(79.8)=-62.5$, $p<0.001$). (The table containing the distances between all relevant language pairs is available on OSF: https://osf.io/3he75/.)

**Text S1: Robustness of the results to the materials used (Bible vs. Alice).**

As described in Methods, two sets of materials were used: one set of materials came from the publicly available corpus of Bible audio stories (n=18), and the second set consisted of passages from Alice in Wonderland (n=16). To test whether the choice of materials affected the pattern of condition responses in the language network, we fit the following LME model with Materials as a fixed effect:

$$EffectSize \sim Materials + (1|Condition) + (1|Participant) + (1|ROI)$$

The results indicated that the overall condition pattern in the language network was robust to the materials used: the Materials effect was not reliable ($\beta$=-0.137, $p$=0.603) (cf. Figures S3 and S5.)

**Text S2: Robustness of the results to participants' age.**

As described in Methods, the age of the participants ranged from 19 to 71 years at the time of testing (*M*=35.0 years, *SD*=13.9). To test whether age affected the pattern of condition responses in the language network, we fit the following LME model with Age as a fixed effect:

$$EffectSize \sim Age + (1|Condition) + (1|Participant) + (1|ROI)$$

The results indicated that the overall condition pattern in the language network was robust to the participants' age: the Age effect was not reliable (β=-0.005, p=0.617).

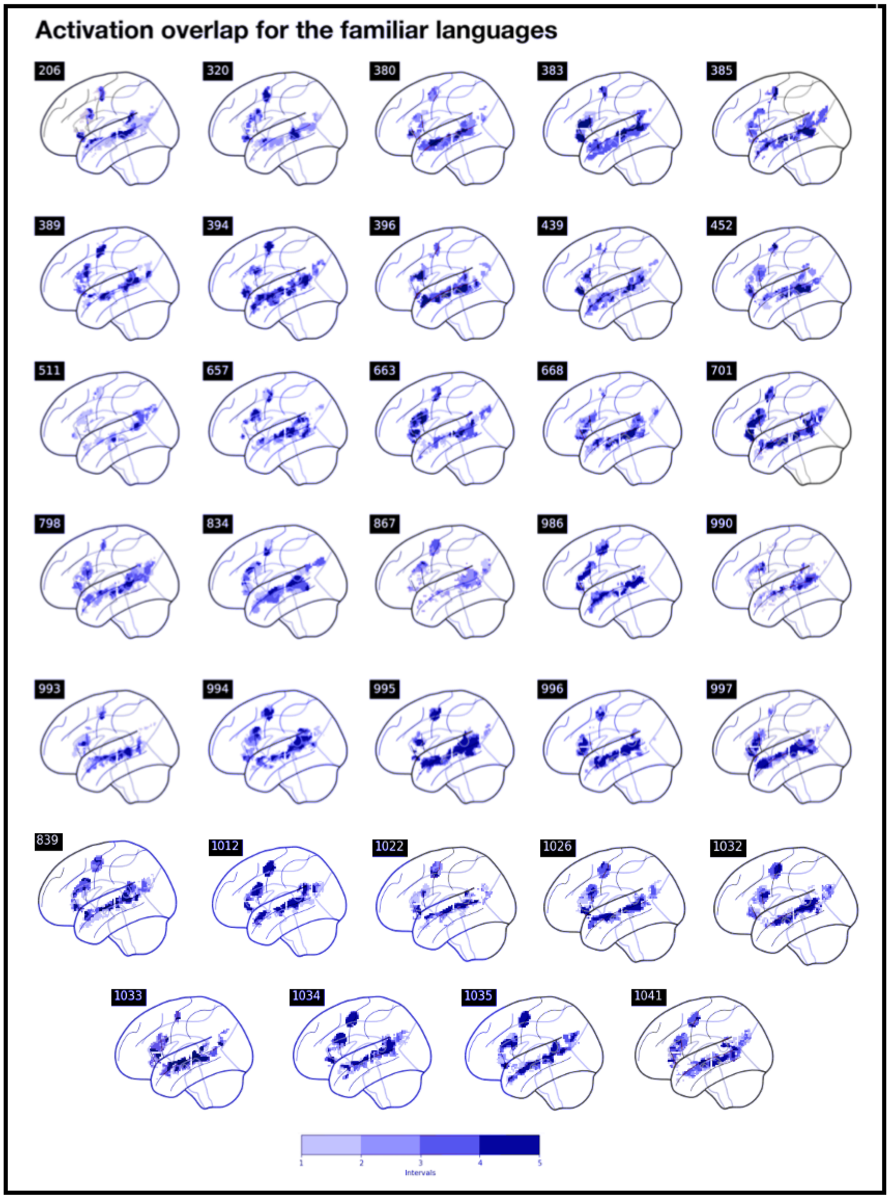


**Figure S1: Activation overlap for the familiar languages (L1-L4) in all participants (n=34).** For each language, we selected 10% of voxels in the left hemisphere that were most responsive to the Language > Quilted-control contrast (based on the contrast values). The activations are shown within the boundaries of the language parcels (see Methods). Colors correspond to the *number of languages* (between 1 and 4) for which the voxel was in the set of top 10% of most responsive voxels. (Three-digit numbers in black boxes correspond to the unique ID of the participant and can be cross-referenced with the data on OSF: https://osf.io/3he75/.)

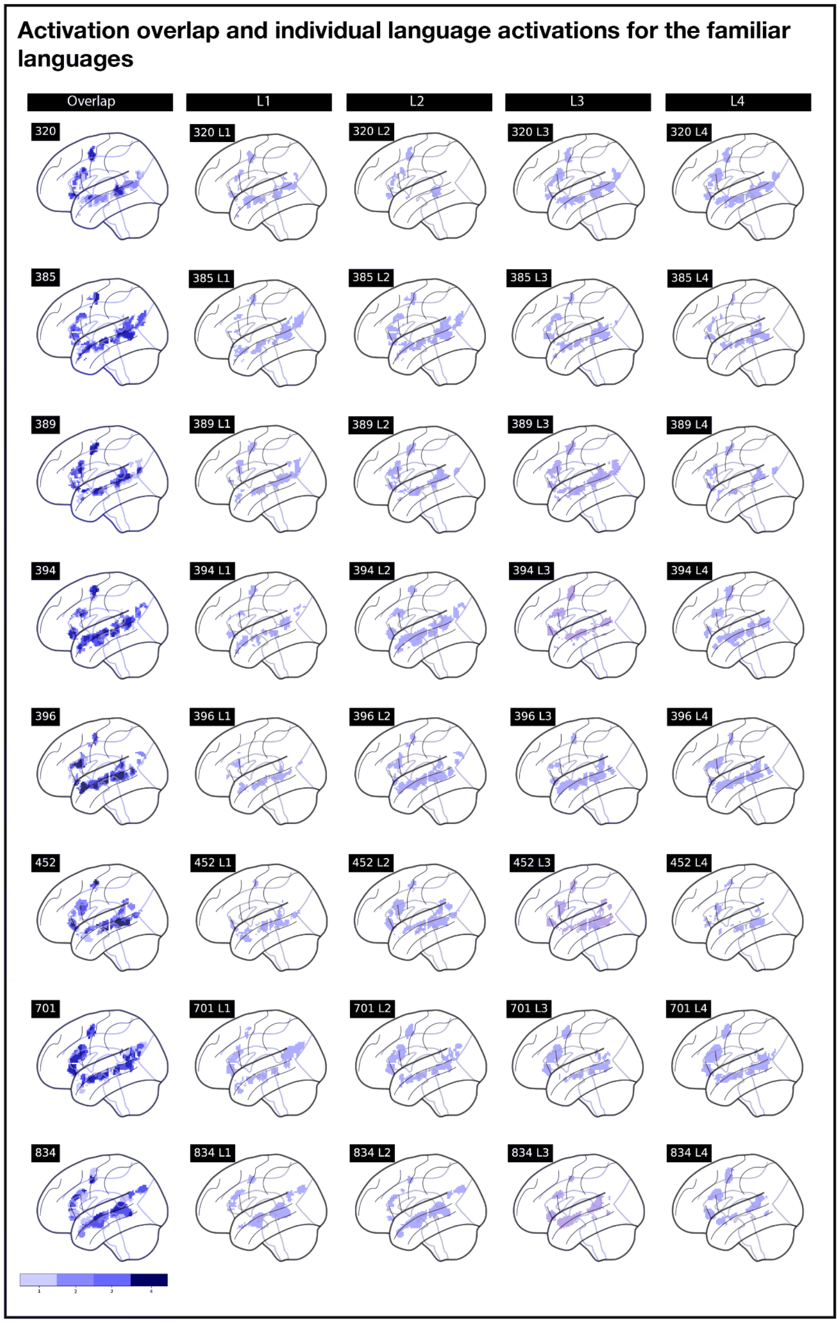


**Figure S2: Activation overlap and individual language activations for the familiar languages (L1-L4).** For each language, we selected 10% of voxels in the left hemisphere that were most responsive to the Language > Quilted-control contrast (based on the contrast values). The activations are shown within the boundaries of the language parcels (see Methods). In the first column, colors correspond to the *number of languages* (between 1 and 4) for which the voxel was in the set of top 10% of most responsive voxels. In the next three columns, activation maps for L1, L2, L3, and L4 are shown. (Three-digit numbers in black boxes correspond to the unique ID of the participant and can be cross-referenced with the data on OSF: https://osf.io/3he75/. Depicted here are the eight polyglots that are included in Figure 1a in the main text; the overlap maps and the individual language activation maps for all polyglots can be found on OSF.)

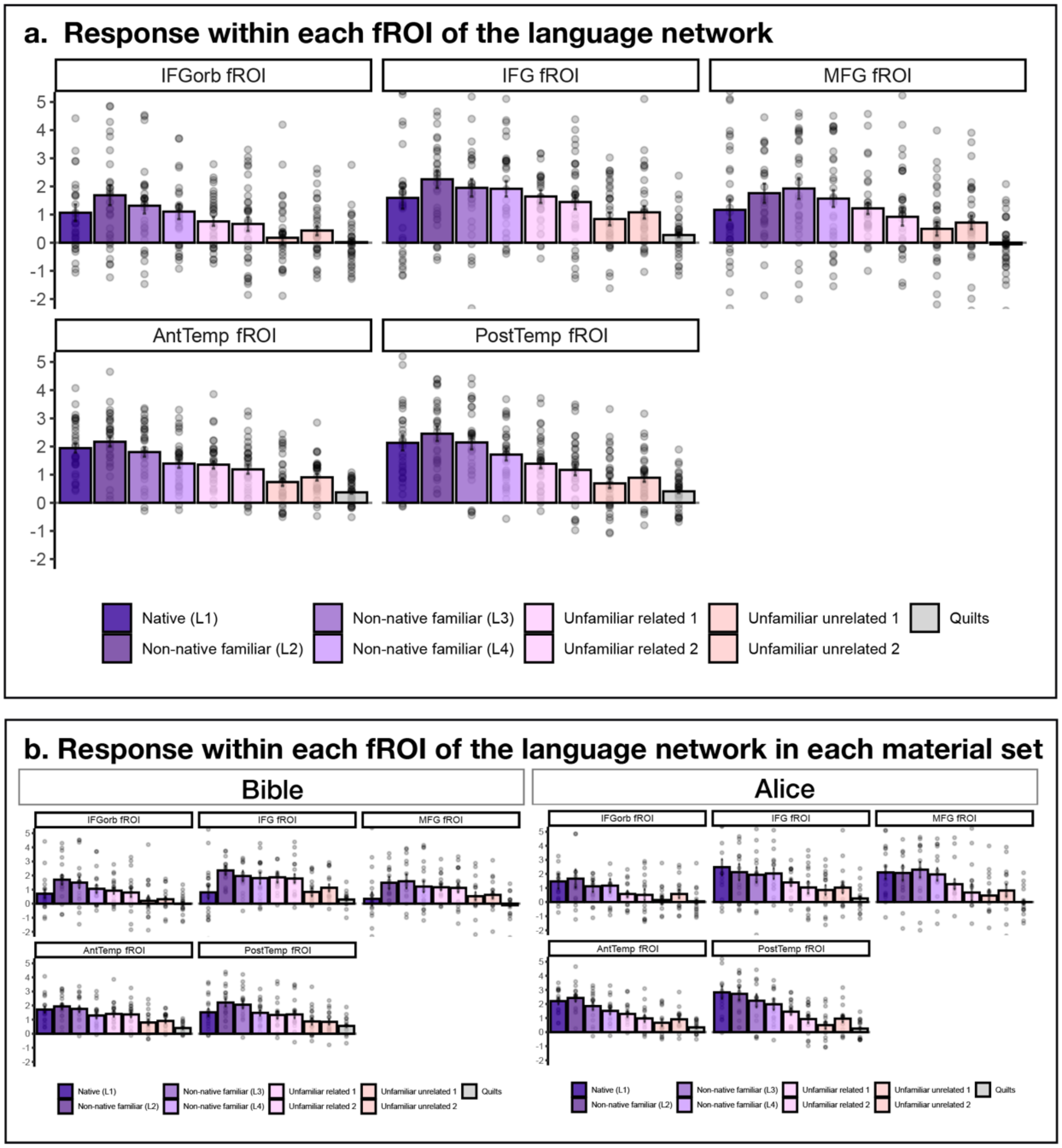


**Figure S3**: **Responses to different languages in the polyglots' language network shown for each fROI separately, including broken up by the materials used (Bible vs. Alice).** Response in the language fROIs to the conditions of the multi-language experiment relative to the fixation baseline (panel **a**: the responses across the full set of 34 participants; panel **b**: participants are split by the version of the experiment). The conditions include the participant's native language (L1), three non-native languages that the participant is somewhat proficient in (L2, L3, and L4; proficiency is highest for L2, lower for L3, and lower still for L4, as described in Methods), four unfamiliar languages (two languages that are related to the languages that the participant is relatively proficient in, two languages that the participant is completely unfamiliar with), and the perceptually matched control condition (Quilts; see Methods). Dots correspond to individual participants, error bars represent standard errors of the mean by participant. The language fROIs are defined by the Sentences > Nonwords contrast in the English localizer (see Methods; see Figure S4 for evidence that the results are similar when the L1 > Quilted contrast from the critical task is used as the localizer).

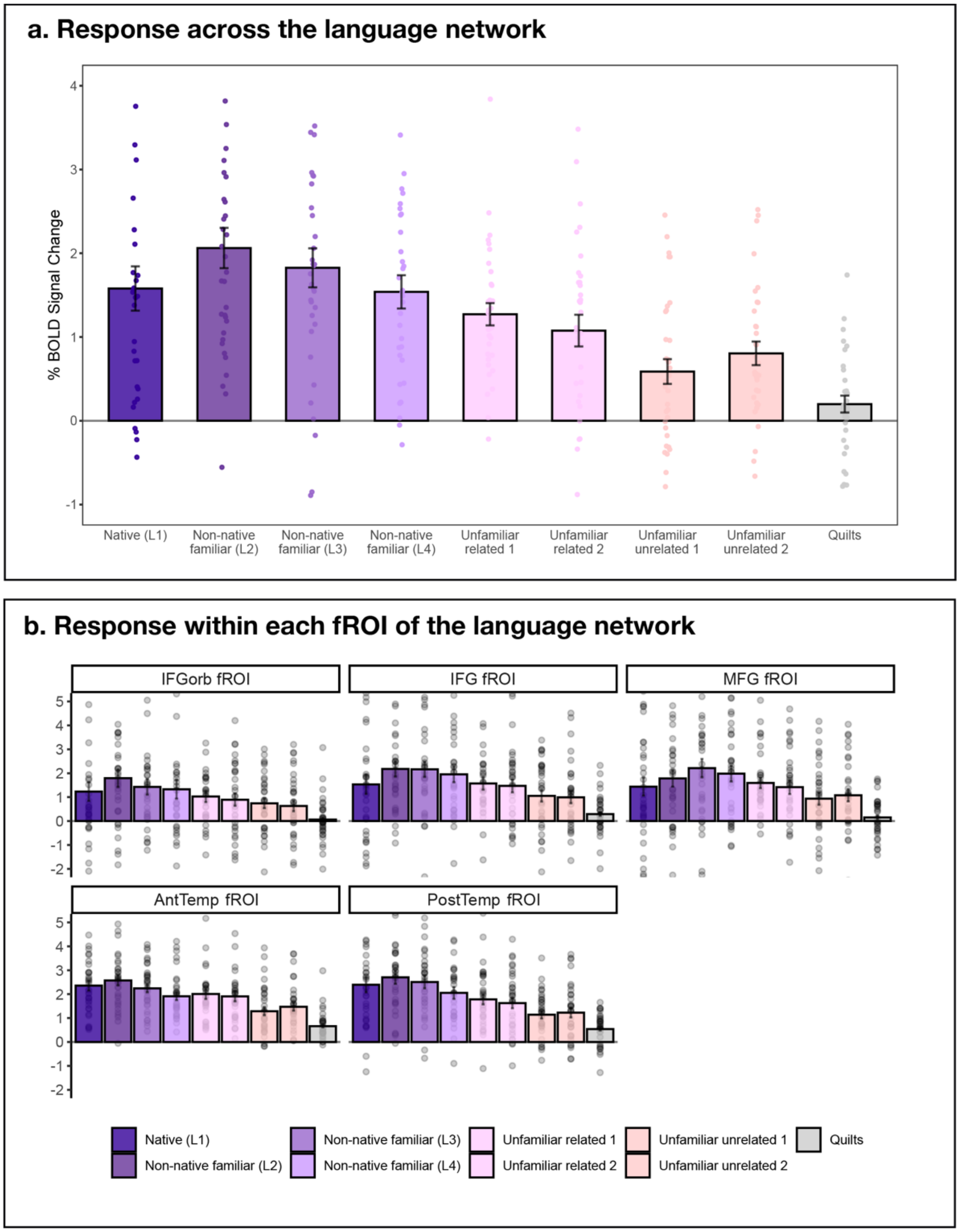


**Figure S4**: **Responses to different languages in the polyglots' language network (defined by the L1 > Quilted contrast).** Response in the language network to the conditions of the multi-language experiment relative to the fixation baseline (panel **a**: the responses averaged across the five fROIs; panel **b**: the responses shown for each fROI separately). The conditions include the participant's native language (L1), three non-native languages that the participant is somewhat proficient in (L2, L3, and L4; proficiency is highest for L2, lower for L3, and lower still for L4, as described in Methods), four unfamiliar languages (two languages that are related to the languages that the participant is relatively proficient in, two languages that the participant is completely unfamiliar with), and the perceptually matched control condition (Quilts; see Methods). Dots correspond to individual participants, error bars represent standard errors of the mean by participant.

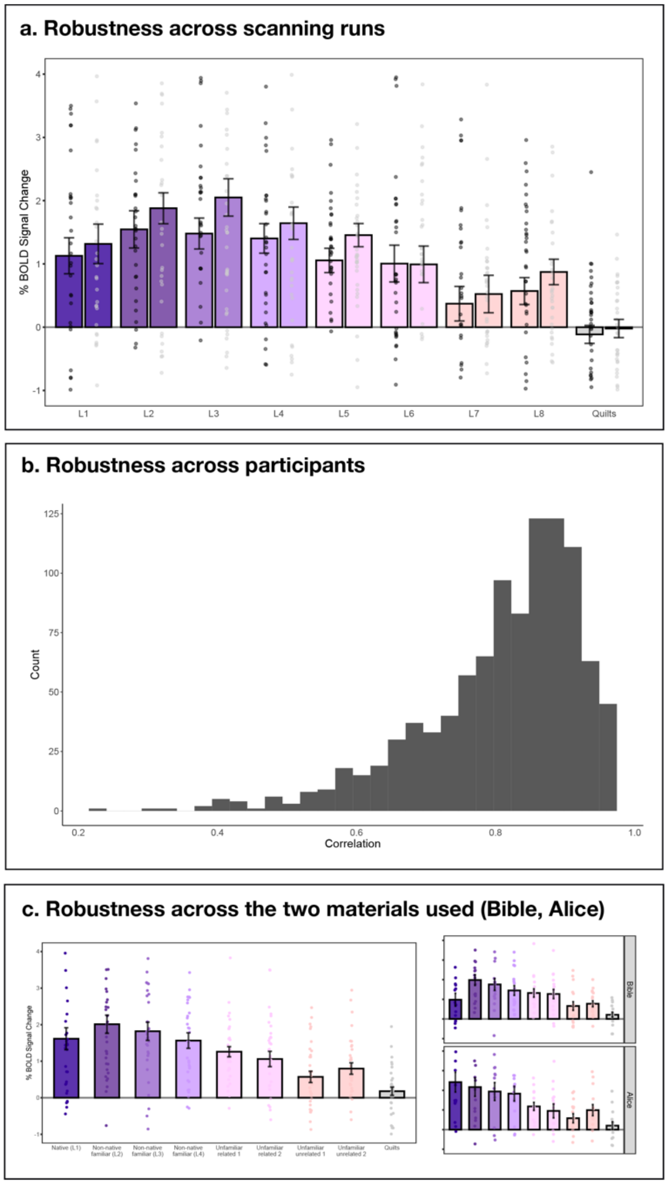


**Figure S5: Robustness of the results across runs, participants, and the materials used (Bible vs. Alice). a.** Robustness of the results across scanning runs. Response in the language network to the conditions of the multi-language experiment relative to the fixation baseline broken down by odd- vs. even-numbered runs. Here and in panel c, dots correspond to individual participants, error bars represent standard errors of the mean by participant. **b.** Robustness of the results across participants. To estimate the robustness of the response profile across participants, we performed an analysis where we divided the set of 34 participants into two random subsets (of size n=17) and computed a correlation for the responses to the nine conditions across the two subsets. We performed this procedure 1,000 times. The figure shows the distribution of the 1,000 resulting correlation values (*M*=0.89, *SD*=0.65, range: 0.57-0.99). **c.** Robustness of the results across the materials used (Bible vs. Alice). As described in Methods, two sets of materials were used: one set of materials came from the publicly available corpus of Bible audio stories (n=18; shown in the top panel), and the second set consisted of passages from Alice in Wonderland (n=16; shown in the bottom panel). The larger panel shows the pattern across all participants (same as main Figure 1b).

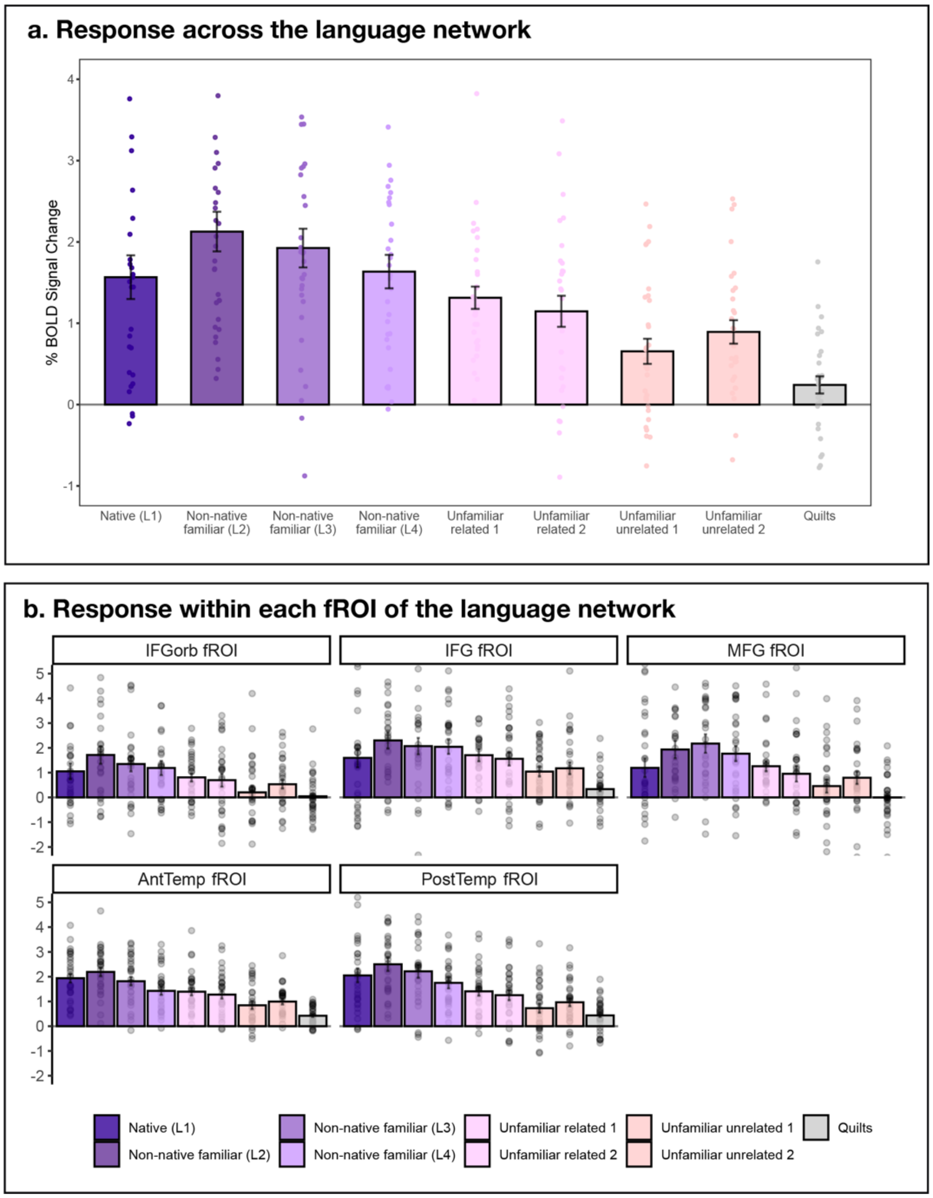


**Figure S6: Responses to different languages in the polyglots' language network when excluding participants whose native language proficiency is below the maximum score of 20 (participant UIDs: 511, 839, 1033).** Response in the language network to the conditions of the multi-language experiment relative to the fixation baseline (panel **a**: the responses averaged across the five fROIs; panel **b**: the responses shown for each fROI separately). The conditions include the participant's native language (L1), three non-native languages that the participant is somewhat proficient in (L2, L3, and L4; proficiency is highest for L2, lower for L3, and lower still for L4, as described in Methods), four unfamiliar languages (two languages that are related to the languages that the participant is relatively proficient in, two languages that the participant is completely unfamiliar with), and the perceptually matched control condition (Quilts; see Methods). Dots correspond to individual participants, error bars represent standard errors of the mean by participant.

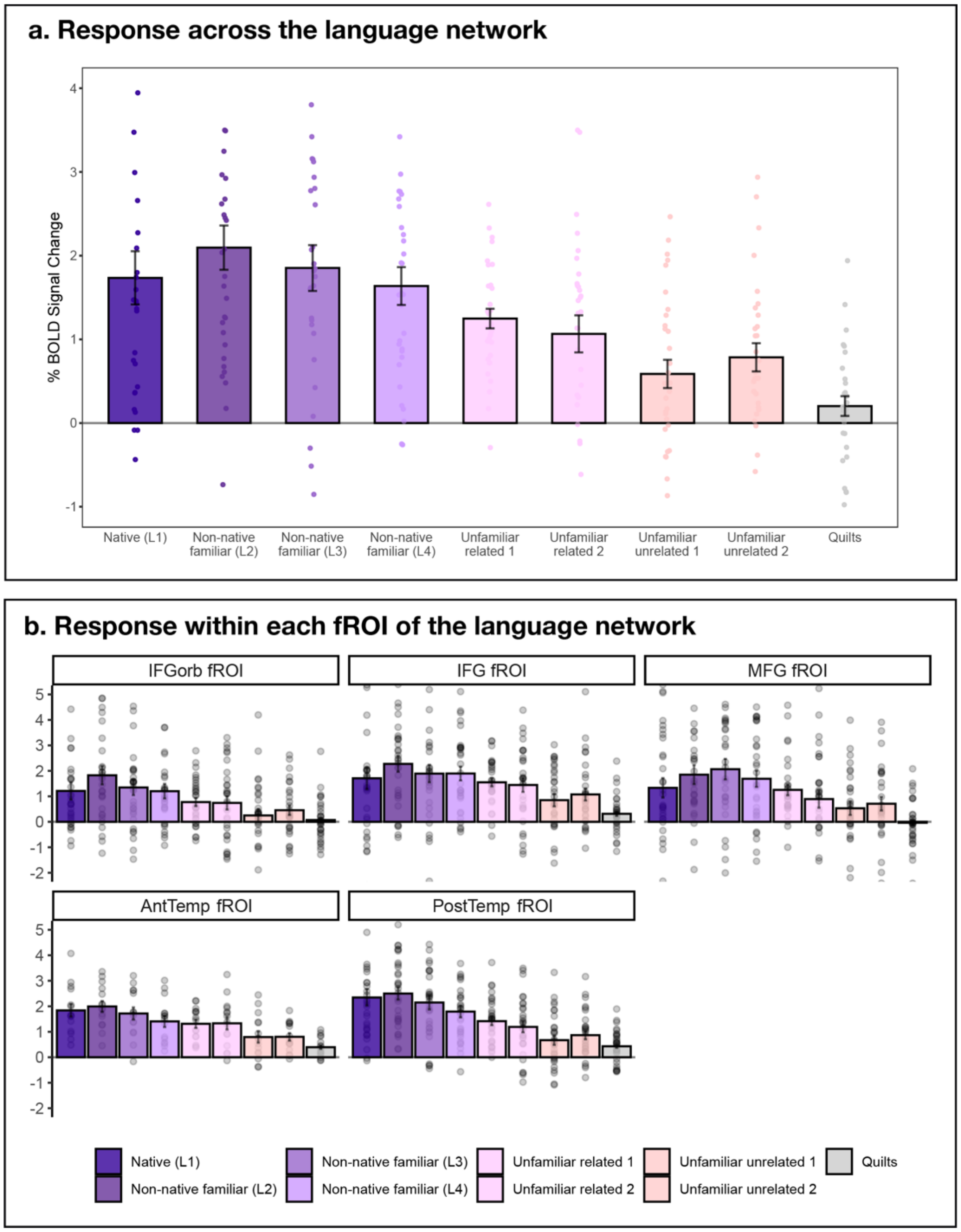


**Figure S7: Responses to different languages in the polyglots' language network when excluding participants whose L2 is also a native language (i.e., early balanced bilinguals; participant UIDs: 383, 798 and 867).** Response in the language network to the conditions of the multi-language experiment relative to the fixation baseline (panel **a**: the responses averaged across the five fROIs; panel **b**: the responses shown for each fROI separately). The conditions include the participant's native language (L1), three non-native languages that the participant is somewhat proficient in (L2, L3, and L4; proficiency is highest for L2, lower for L3, and lower still for L4, as described in Methods), four unfamiliar languages (two languages that are related to the languages that the participant is relatively proficient in, two languages that the participant is completely unfamiliar with,) and the perceptually matched control condition (Quilts; see Methods). Dots correspond to individual participants, error bars represent standard errors of the mean by participant.

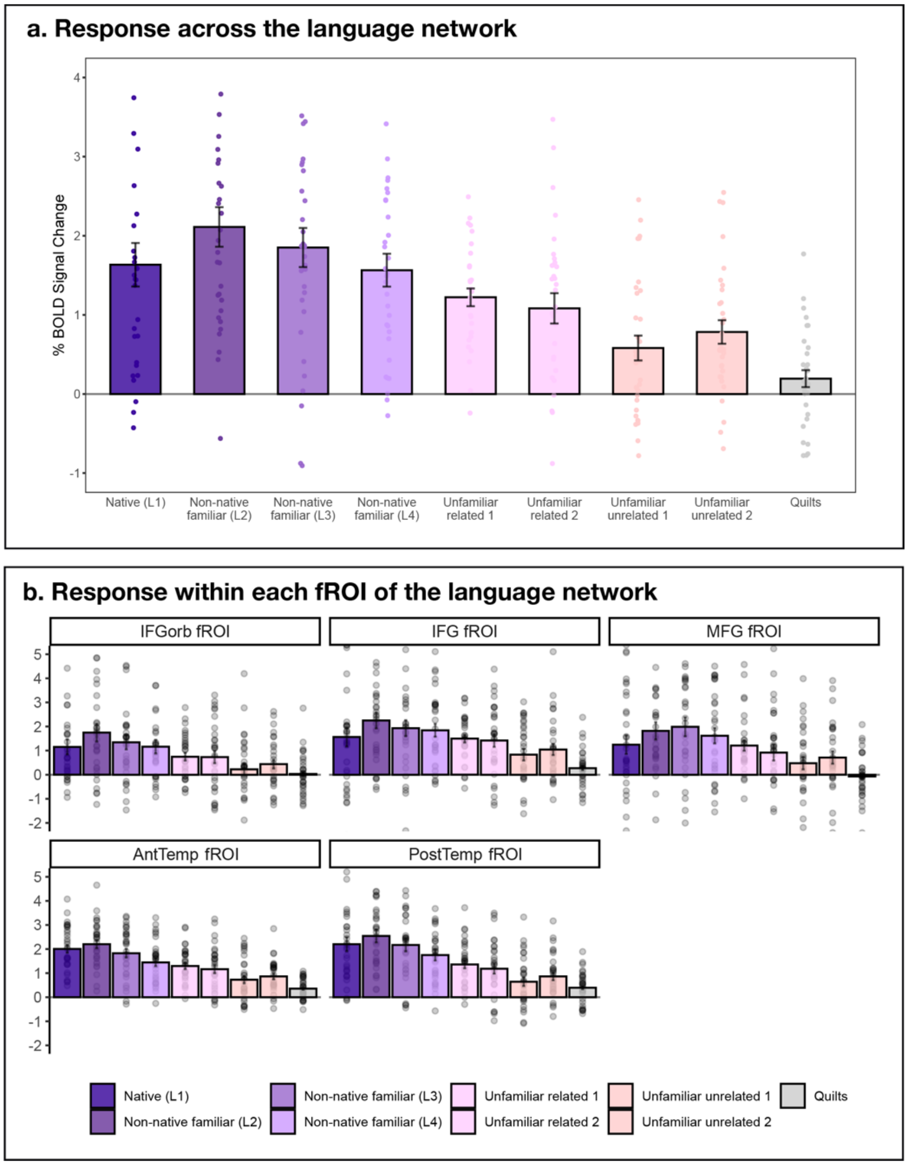


**Figure S8: Responses to different languages in the polyglots' language network when excluding participants with errors in the selection of related languages (participant UIDs: 383 and 867).** Response in the language network to the conditions of the multi-language experiment relative to the fixation baseline (panel **a**: the responses averaged across the five fROIs; panel **b**: the responses shown for each fROI separately). The conditions include the participant's native language (L1), three non-native languages that the participant is somewhat proficient in (L2, L3, and L4; proficiency is highest for L2, lower for L3, and lower still for L4, as described in Methods), four unfamiliar languages (two languages that are related to the languages that the participant is relatively proficient in, two languages that the participant is completely unfamiliar with), and the perceptually matched control condition (Quilts; see Methods). Dots correspond to individual participants, error bars represent standard errors of the mean by participant.

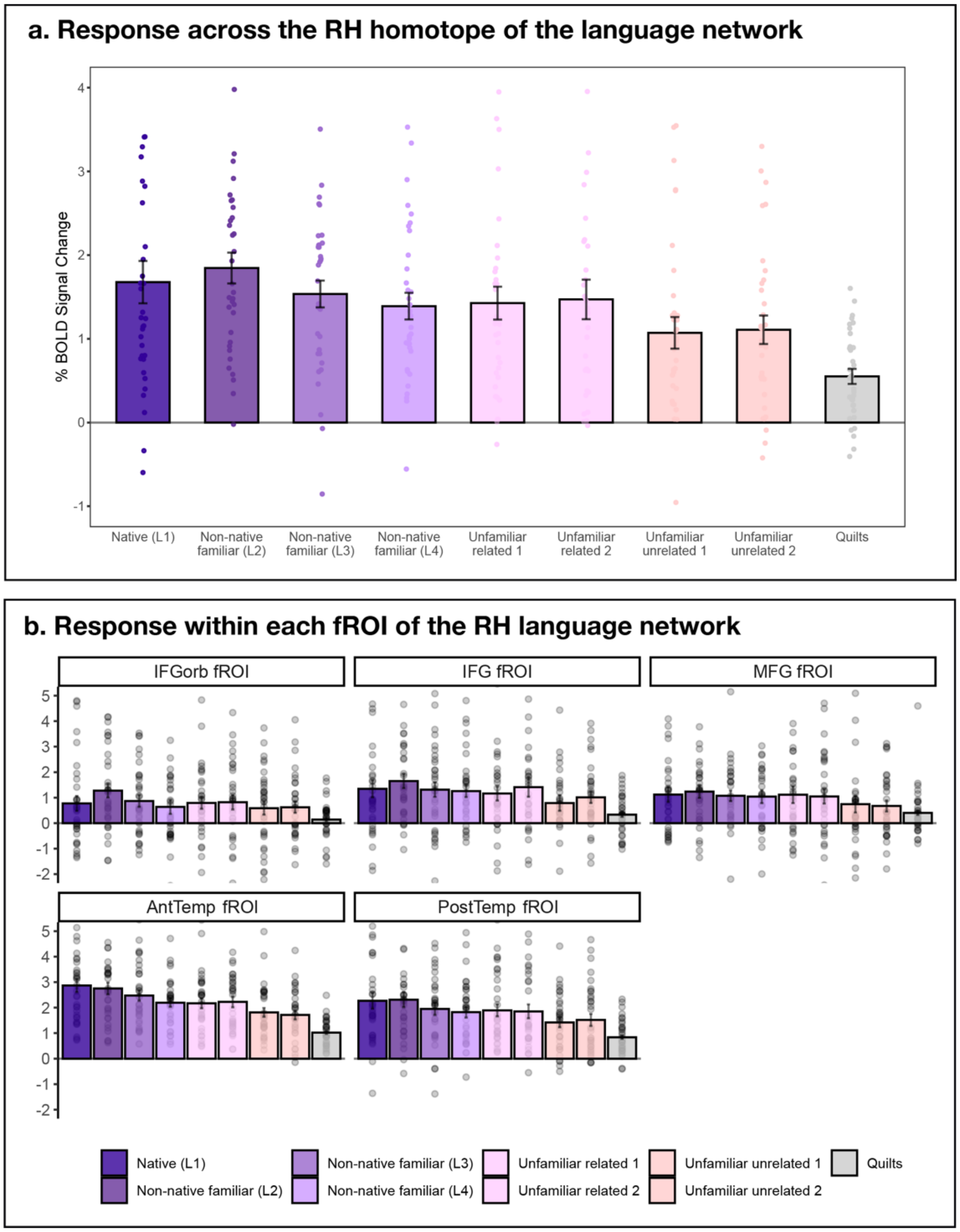


**Figure S9: Responses to different languages in the polyglots' right hemisphere homotope of the language network.** Response in the language network to the conditions of the multi-language experiment relative to the fixation baseline (panel **a**: the responses averaged across the five fROIs; panel **b**: the responses shown for each fROI separately). The conditions include the participant's native language (L1), three non-native languages that the participant is somewhat proficient in (L2, L3, and L4; proficiency is highest for L2, lower for L3, and lower still for L4, as described in Methods), four unfamiliar languages (two languages that are related to the languages that the participant is relatively proficient in, two languages that the participant is completely unfamiliar with), and the perceptually matched control condition (Quilts; see Methods). Dots correspond to individual participants, error bars represent standard errors of the mean by participant. Similar to the LH language network (main text Figure 1b), all languages elicited a reliable response relative to the control condition (*p*s<0.001). However, the difference between non-native familiar (L2, L3, L4) and unfamiliar languages was not as pronounced in the RH homotopic areas, as revealed by a significant Hemisphere by Condition interaction: $p=0.010$ (in the LH, the mean response was 1.81 for familiar languages and 0.95 for unfamiliar languages, whereas in the RH, the mean response was 1.45 for familiar languages and 1.04 for unfamiliar languages).

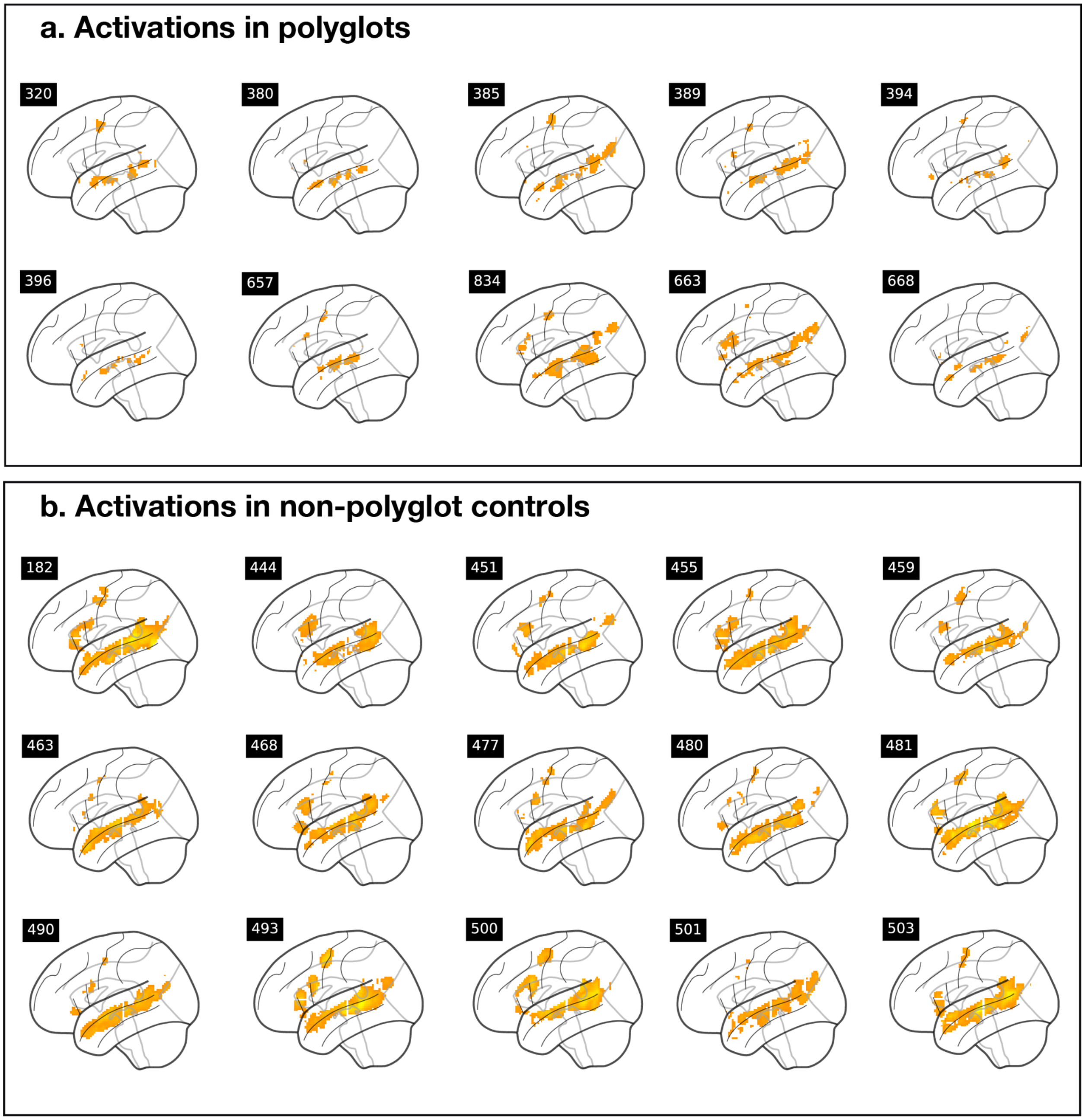


**Figure S10: Activation maps for polyglots and non-polyglot control participants for native language processing.** For each participant, we selected voxels that are significant (at the p<0.001 uncorrected whole-brain threshold) for the L1 > Quilted control contrast. The activations are shown within the boundaries of the language parcels (see Methods). In line with the lower magnitude of response in the individually defined language fROIs for polyglots compared to non-polyglots (Figure 2 in the main text), activations are less spatially extensive for polyglots than non-polyglots at a fixed threshold. (Three-digit numbers in black boxes correspond to the unique ID of the participant and can be cross-referenced with the data on OSF: https://osf.io/3he75/. Depicted here are subsets of participants in each group; the maps for all participants (n=34 polyglots and n=86 non-polyglots) can be found on OSF: https://osf.io/3he75/.)

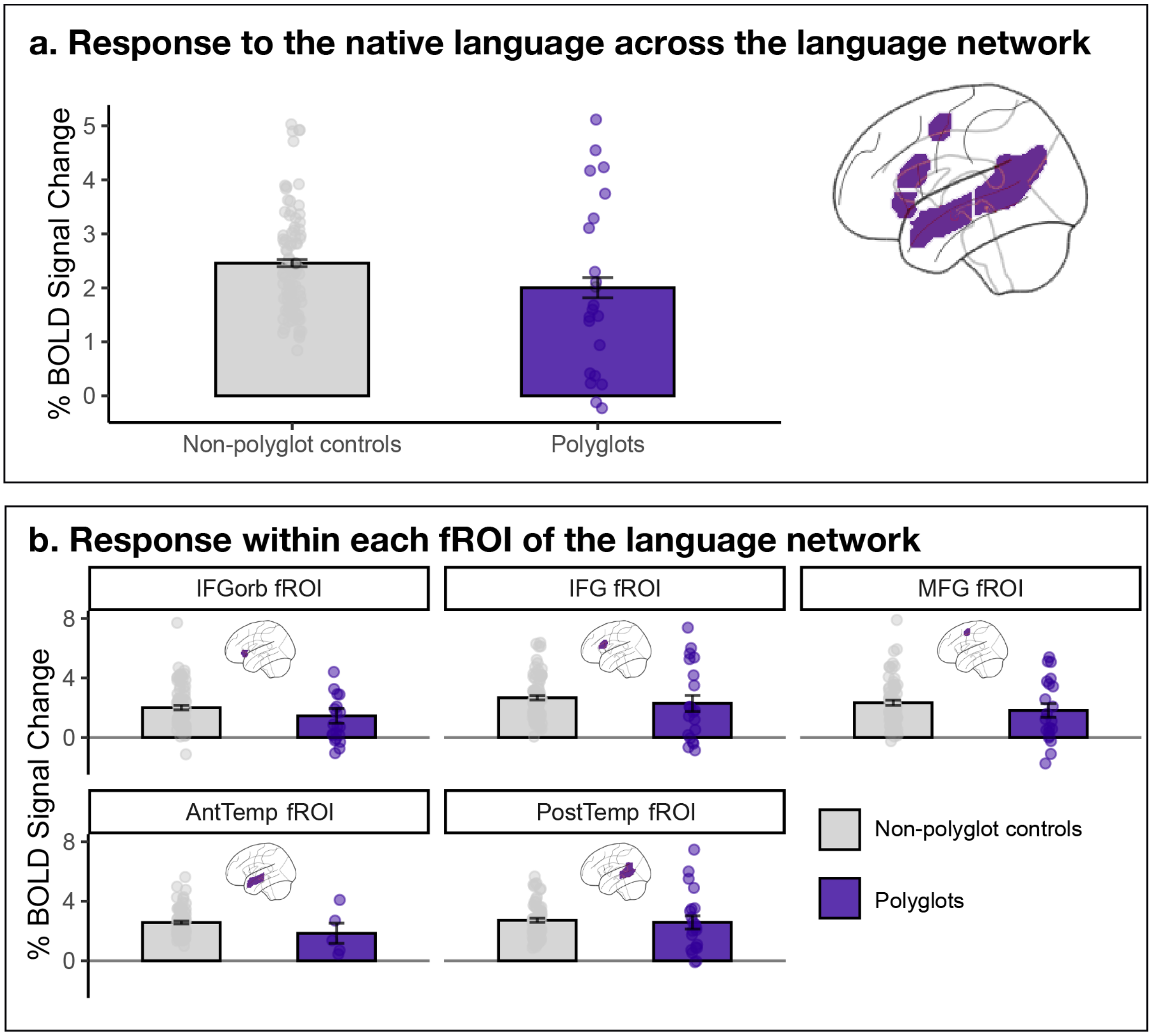


**Figure S11: Response to the native language in the language network of polyglots (purple, n=19 participants who were not included in Jouravlev et al., 2021) and non-polyglot controls (grey, n=86).** Response in the language network to the native language condition of the multi-language experiment relative to the fixation baseline (panel **a**: the responses averaged across the five fROIs; panel **b**: the responses shown for each fROI separately). Polyglots showed a numerically lower response in their language network while listening to their native language than a control group of 86 bilingual non-polyglots (1.79 vs. 2.46 % BOLD signal change relative to the fixation baseline; β=-0.53, p=0.063). Dots correspond to individual participants, error bars represent standard errors of the mean by participant. (The three participants who rated their native language proficiency lower than the maximum of 20 were excluded from this analysis.)

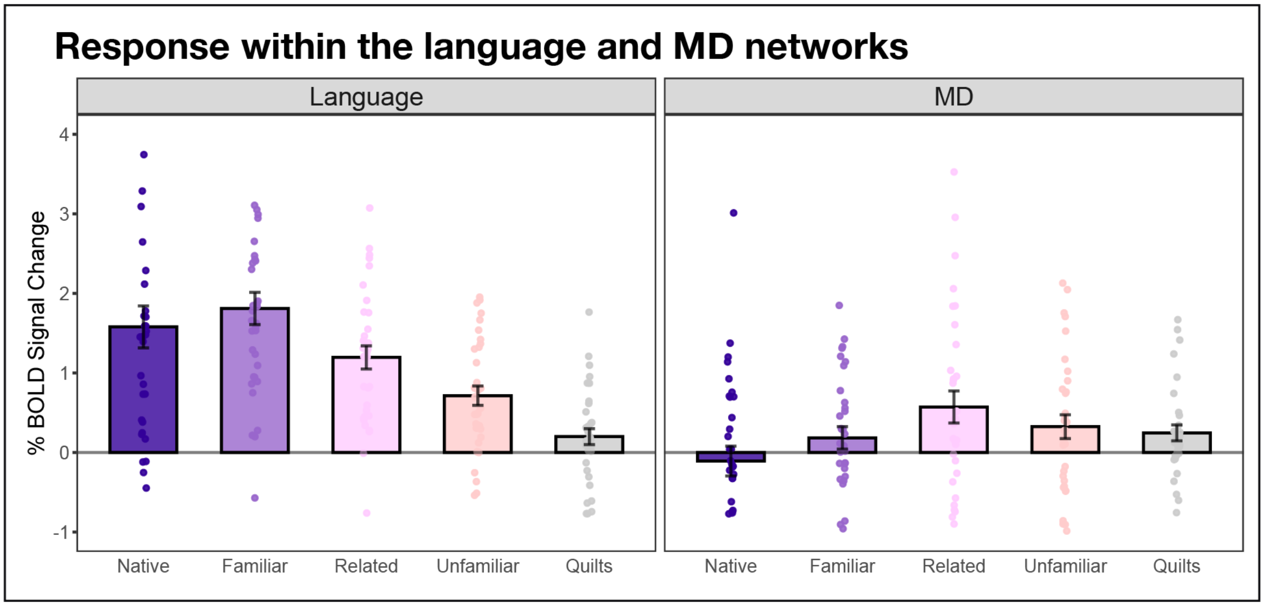


**Figure S12: Responses to different language conditions in the polyglots' language and Multiple Demand (MD) networks.** Response in the language network (averaging across the five fROIs) and in the Multiple Demand (MD) network (averaging across 20 fROIs; e.g., Malik-Moraleda, Ayyash et al., 2022) to the condition types in the multi-language experiment relative to the fixation baseline. The MD fROIs were defined by a Hard > Easy contrast in a spatial working memory task (where participants keep track of more vs. fewer locations in a grid; Fedorenko et al., 2013), which robustly identifies the MD network at the individual-participant level (e.g., Blank et al., 2014; Shashidhara et al., 2019; Assem et al., 2020b; Malik-Moraleda et al., 2021). The conditions include the participant's native language (Native), three familiar non-native languages (L2-L4) that the participant is somewhat proficient in (Familiar), two unfamiliar languages that are related to the languages that the participant is relatively proficient in (Related), two languages that the participant is completely unfamiliar with (Unrelated), and the perceptually matched control condition (Quilts; see Methods). Dots correspond to individual participants, error bars represent standard errors of the mean by participant. In line with much past work (e.g., Malik-Moraleda, Ayyash et al., 2022), the responses to auditory language comprehension in the MD network are generally low. Importantly, however, familiar, related, and unrelated languages (as well as the Quits control condition) all elicit an above-baseline response (albeit a weak one; the Related condition elicits the highest response), but native language processing elicits, on average, a response below the fixation baseline.

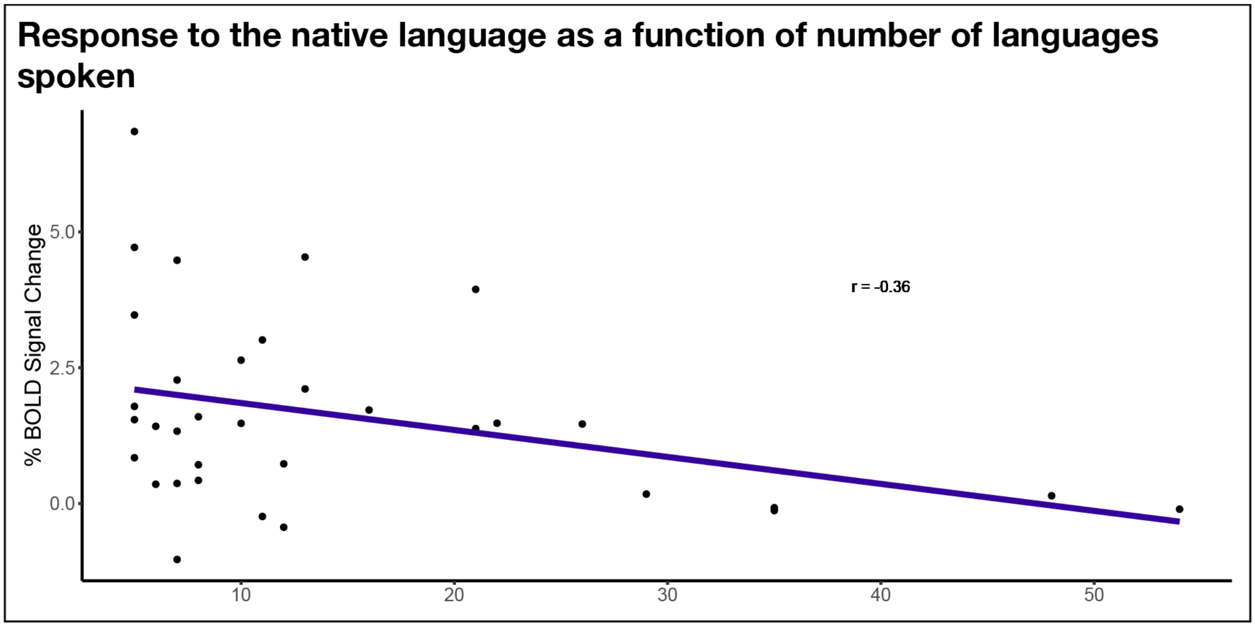

**Figure S13: Response to the native language as a function of the number of languages spoken by the polyglot.**

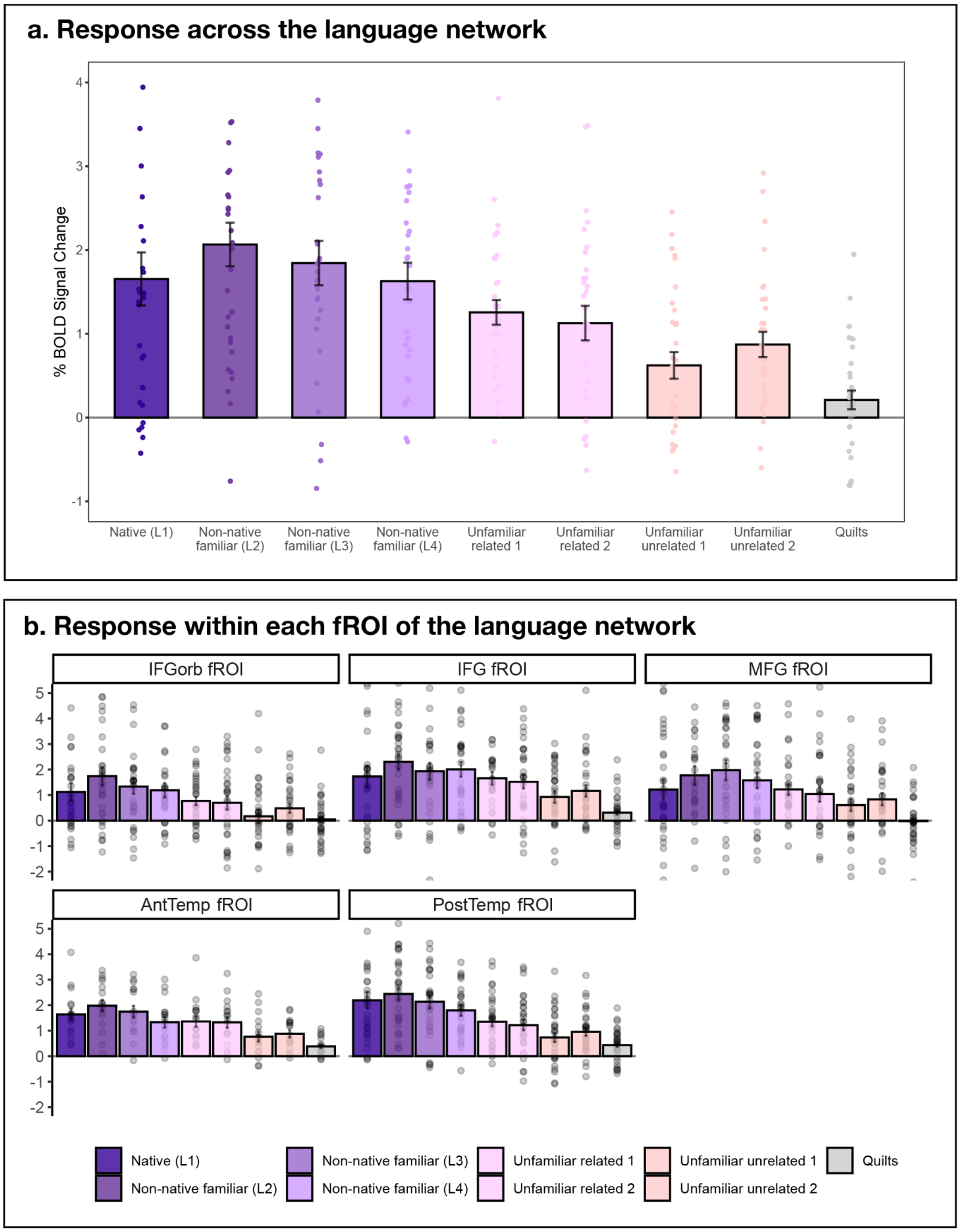


**Figure S14: Responses to different languages in the polyglots' language network when excluding left-handed participants (n=2).** Response in the language network to the conditions of the multi-language experiment relative to the fixation baseline (panel **a**: the responses averaged across the five fROIs; panel **b**: the responses shown for each fROI separately). The conditions include the participant's native language (L1), three non-native languages that the participant is somewhat proficient in (L2, L3, and L4; proficiency is highest for L2, lower for L3, and lower still for L4, as described in Methods), four unfamiliar languages (two languages that are related of the languages that the participant is relatively proficient in, two languages that the participant is completely unfamiliar with), and the perceptually matched control condition (Quilts; see Methods). Dots correspond to individual participants, error bars represent standard errors of the mean by participant.

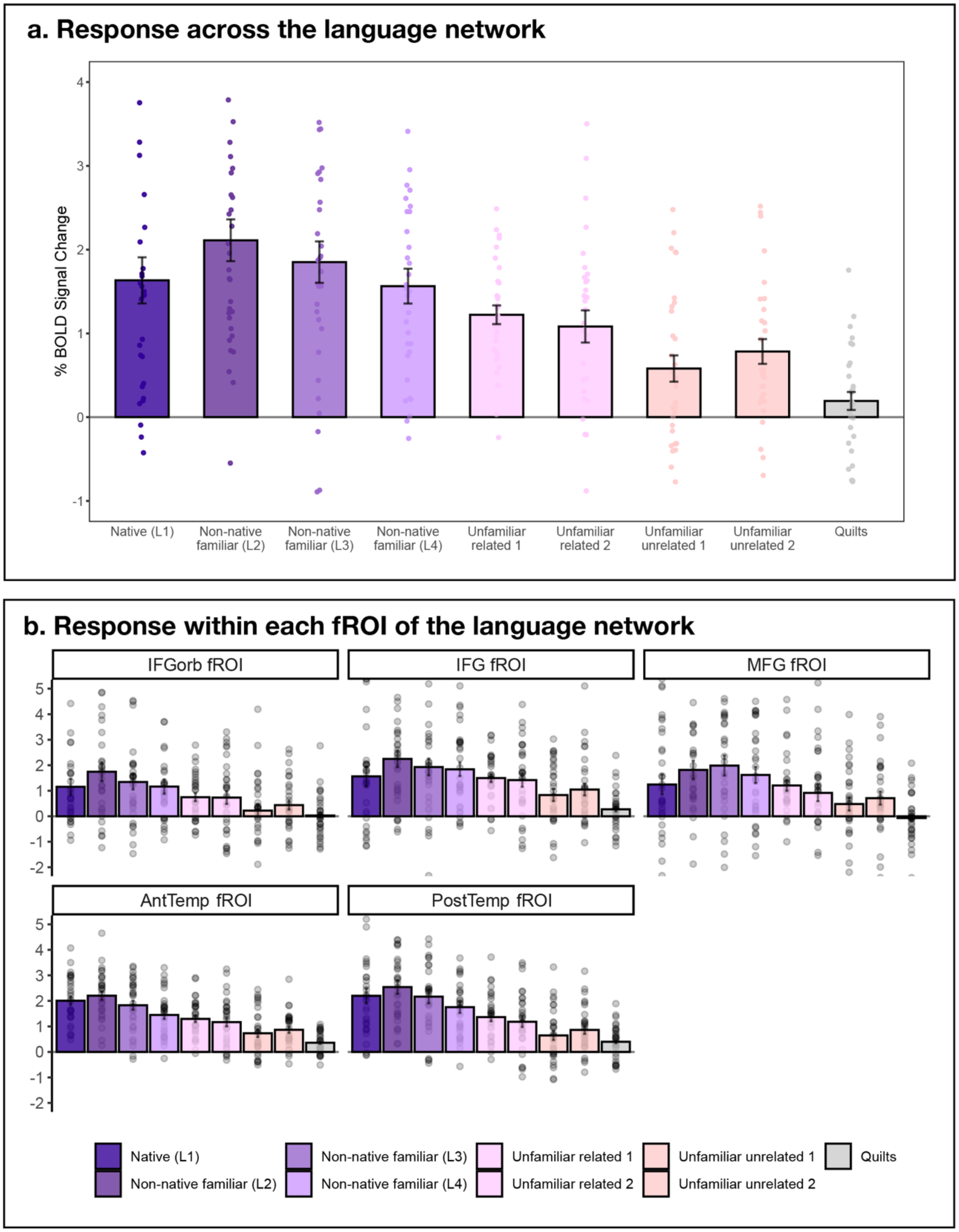


**Figure S15: Responses to different languages in the polyglots' language network in the language-dominant hemisphere.** For one participant (UID 511), the language network was lateralized to the right hemisphere (LI = -0.854; Table S2), and in this version of the analysis, we used the RH language fROIs for this participant. Response in the language network to the conditions of the multi-language experiment relative to the fixation baseline (panel **a**: the responses averaged across the five fROIs; panel **b**: the responses shown for each fROI separately). The conditions include the participant's native language (L1), three non-native languages that the participant is somewhat proficient in (L2, L3, and L4; proficiency is highest for L2, lower for L3, and lower still for L4, as described in Methods), four unfamiliar languages (two languages that are related of the languages that the participant is relatively proficient in, two languages that the participant is completely unfamiliar with), and the perceptually matched control condition (Quilts; see Methods). Dots correspond to individual participants, error bars represent standard errors of the mean by participant.
